# A dynamic *in vitro* model of Down Syndrome neurogenesis with Trisomy 21 gene dosage correction

**DOI:** 10.1101/2022.05.11.491519

**Authors:** Prakhar Bansal, Erin. C Banda, Heather R. Glatt-Deeley, Christopher E. Stoddard, Jeremy W. Linsley, Neha Arora, Darcy T. Ahern, Yuvabharath Kondaveeti, Michael Nicouleau, Miguel Sabariego-Navarro, Mara Dierssen, Steven Finkbeiner, Stefan F. Pinter

## Abstract

Excess gene dosage from human chromosome 21 (chr21) causes Down syndrome (DS), spanning developmental as well as acute phenotypes in terminal cell types. Which phenotypes remain amenable to intervention after development is unknown. To address this question in a model of DS neurogenesis, we generated trisomy 21 (T21) human induced pluripotent stem cells (hiPSCs) alongside otherwise isogenic euploid controls from mosaic DS fibroblasts, and integrated an inducible *XIST* transgene on one chr21 copy. Monoallelic chr21 silencing by *XIST* was near-complete and irreversible in hiPSCs. Differential expression reveals T21 neural lineages and T21 hiPSCs suppress similar translation and mitochondrial pathways, and activate cellular stress responses. When *XIST* is induced before the neural progenitor stage, T21 dosage correction mitigates a pronounced skew towards astrogenesis in differentiation. Because our transgene remained inducible in post-mitotic T21 neurons and astrocytes, we demonstrate *XIST* efficiently represses genes even after terminal differentiation, which will empower exploration of cell type-specific T21 phenotypes that remain responsive to chr21 dosage.

## INTRODUCTION

As the most prevalent human viable aneuploidy (trisomy 21, T21 in up to 1:800-1:1000 live births), Down syndrome (DS) poses three significant challenges to understanding and addressing the clinical needs of people living with DS ^1^. First, our understanding of the developmental etiology underlying cardiac, hematopoietic and neurological defects in DS is still limited ^2^. Ever-more sophisticated rodent DS models faithfully carry most of human chromosome 21 (chr21) ^3,4^ or traverse syntenic boundaries to represent supernumerary mouse chr21 orthologs ^5^. While these rodent models capture many important aspects of DS, human T21 induced pluripotent stem cells (hiPSCs) are serving as an important complementary model for DS development and pre-clinical research ^6^. These hiPSC models may prove especially relevant to human-specific aspects of development, for example by reflecting brain region-specific neural cell types ^7^, intrinsically longer human neuronal maturation ^8^, and a more rapidly evolving susceptibility to neurodegeneration that may be rooted in human neurodevelopment ^9,10^.

Second, we lack experimental approaches to systematically distinguish irreversible developmental phenotypes from “acute” cellular phenotypes that may remain amenable to postnatal intervention. Supernumerary chr21 genes, or their rodent orthologs, are present throughout differentiation or development of such DS hiPSC and rodent DS models, respectively. With few exceptions ^11^, we therefore have an incomplete view of which T21 phenotypes depend on ongoing excess dosage of specific chr21 genes in terminally differentiated cells. This question is particularly relevant to DS-associated neurodegeneration ^12^.

Third, cellular stress pathways that are chronically and systemically active in DS ^13–18^ and likely play an important role in DS neurodegeneration ^19^, are also shared with human trisomies 13 (T13) and 18 (T18) ^20^, but not with dosage-compensated sex-chromosomal trisomy. The most plausible origin for such common aneuploidy-associated cellular stress is the cumulative excess expression of the many autosomal genes that code for subunits of dosage-sensitive protein complexes. However, this interpretation has not yet been tested formally in the context of an autosomal trisomy, as it requires distinguishing excess expression from the mere presence of the supernumerary chromosome.

A human cellular model with dynamic T21 gene dosage would be instrumental towards addressing these challenges, across a range of DS-affected cell fates and phenotypes. To characterize the impact of T21 in hiPSCs and terminally differentiated cell types, we generated our own dynamic hiPSC model of T21 dosage alongside otherwise isogenic controls from mosaic DS. Pioneering work by the Lawrence lab demonstrated the utility of the long non-coding RNA *XIST*, which endogenously silences the inactive X in females, for T21 dosage correction in DS hiPSCs ^21^. *XIST* RNA spreads from its site of transcription across the entire chromosome *in cis*, to directly and indirectly recruit orthogonal repressive chromatin complexes to silence genes ^22^. Yet, earlier work in the mouse system indicated that *Xist* could establish gene silencing only during or upon exiting pluripotency, but not after differentiation ^23,24^, with rare exception ^25,26^. Recent work by the Lawrence lab shows that human *XIST* can still silence genes up to the neural progenitor cell (NPC) stage, but technical limitations hindered *XIST* induction in terminal, post-mitotic human neural cells ^27^.

Here, we leverage phased allelic expression analysis to demonstrate that our new inducible T21-*XIST* design enables uniform, near-complete and -irreversible T21 dosage correction in hiPSCs. *XIST* suppresses T21 *trans*-effects, including cellular stress pathways that are also active in T21 neural lineages. T21 silencing prior to the NPC stage rescues T21-altered neuro-astroglial lineage balance that reflects excess gliogenesis previously observed in DS fetal samples ^28–32^ and rodent models ^33–35^. Because our T21-*XIST* system is also the first to effectively induce *XIST* and repress chr21 dosage in post-mitotic neurons and astrocytes, this new model of dynamic T21 gene dosage will facilitate identification of cellular pathways and cell types that may present promising targets for intervention after completed development.

## RESULTS

### T21 dosage correction by *XIST-*mediated monoallelic silencing in DS hiPSCs

To derive T21 hiPSCs alongside isogenic euploid control lines, we reprogrammed dermal fibroblasts from a male individual with mosaic DS ^36^, and screened the resulting hiPSC clones using copy-number sensitive qPCR [Fig. S1A,B]. Two T21 lines appeared to have a gain on chr17, a frequent karyotypic abnormality primarily observed in human ESCs ^37,38^, but featured only two centromere 17 signals by fluorescent *in-situ* hybridization (FISH) follow-up [Fig. S1C]. We performed array comparative genomic hybridization (aCGH, CytoSNP-850k) to confirm expected karyotypes for fibroblasts, euploid and T21 hiPSCs, and to resolve segmental (17q) gains in the two corresponding clones, which were excluded thereafter [Fig. S1D]. Differential probe intensity analysis of the remaining T21 and euploid hiPSC lines also revealed a large set of heterozygous SNPs with increased representation in the euploid lines spanning chr21, and a smaller focal set that was lost in euploid lines [Fig. S1E]. These results suggested one chr21 copy from each parent is maintained in euploid lines (chr21^M^ & chr21^P^), whereas T21 hiPSCs carry an additional copy (chr21^M2^), likely a maternal chromatid retained via non-disjunction in meiosis II ^39^.

While T21 is stably maintained in hiPSCs after reprogramming ^40–42^, sporadic T21 loss has been reported ^43,44^. We therefore confirmed the mitotic stability of three chr21 copies by two independent approaches. First, three independent signals were observed in 94.5% of T21 hiPSCs by fluorescent *in-situ* hybridization (FISH, *DSCR8* probe), compared to 99% of euploid hiPSCs with two signals [Fig. S1F]. Second, while targeting a *UBC-*promoter driven doxycycline (dox)-responsive trans-activator (rTTA3G) to the AAVS1 safe harbor in *PPP1R12C* (HSA19), we confirmed that all 32/32 independent colonies arising from single cells maintained T21 by copy-number sensitive qPCR for a *DYRK1A* amplicon [Fig. S1G].

We integrated our TET3G-driven *XIST* cDNA transgene on chr21^P^ [Fig. 1A], after re-verifying the karyotypic integrity of rTTA3G clones by aCGH [Fig. S1D]. To this end, we designed Cas9 guides dependent on a chr21^P^-specific protospacer adjacent motif (PAM), targeting an intergenic site well outside of *DYRK1A* to avoid impacting its expression. We proceeded with three independent, PCR-screened T21-*XIST* clones, and tested for *XIST* induction by RNA-FISH, and immuno-fluorescence (IF) staining for ubiquitinated histone 2A lysine 119 (H2AK119ub) and tri-methylated histone 3 lysine 27 (H3K27me3), which follow *XIST/Xist* RNA expression on mouse and human X alike ^22^. Of note, because the original fibroblasts were from a male individual with DS [Fig. S1C], there was no endogenous *XIST* expression, and any *XIST*/H2A119ub/H3K27me3 cloud must arise from the T21-targeted *XIST* transgene on chr21. Indeed, after cessation of acute dox treatment (3 wk withdrawal, w/d) hiPSCs were negative for *XIST*, underscoring the TET3G system is not leaky and turns off when dox is withdrawn, whereas virtually all T21-*XIST* hiPSCs displayed *XIST* clouds and acquired H2A119ub/H3K27me3 signals in the presence of dox [Fig. 1B]. Together, these data illustrate the homogeneity of *XIST* induction, and provide independent confirmation that this third copy of chr21 is faithfully maintained in hiPSCs.

**Figure 1:**
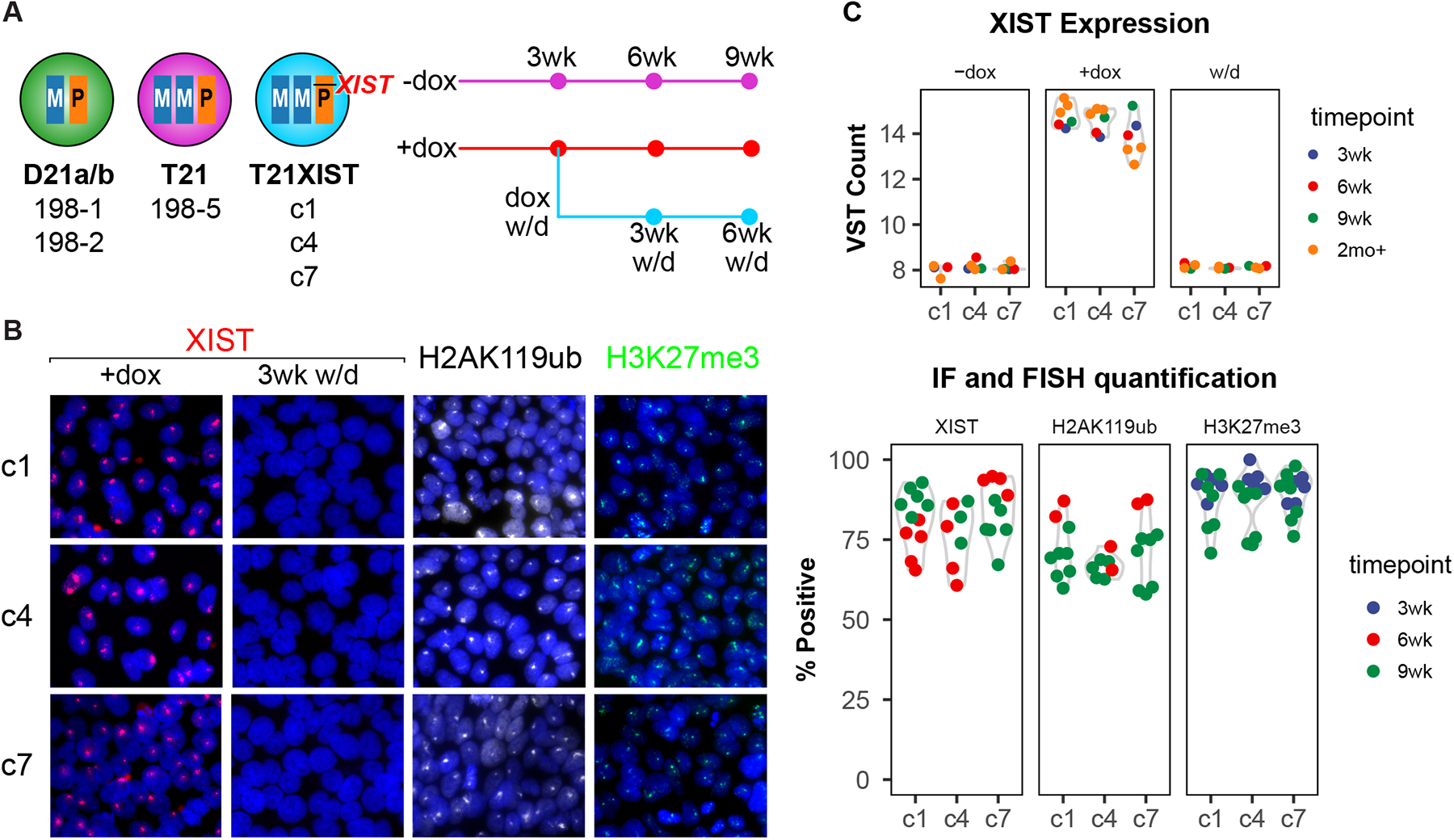
T21 hiPSCs induce *XIST* transgene and deposit heterochromatin on one chr21 allele. (A) Schematic of cell lines and sample collection timepoints and dox treatment conditions. Samples were collected after 3-9 wks of +dox or no-dox treatment, or 3 wk of +dox and 3 or 6 wks of dox withdrawal. A subset of samples was treated with dox for 2mo+ prior to collection/withdrawal. (B) Representative images (left) for XIST-FISH, and H2AK119ub and H3K27me3 IF, and image quantification (right). Control panel of XIST-negative cells after 3wk of dox w/d for XIST-FISH is included on the left. (C) *XIST* expression in T21-XIST samples from mRNA-seq (VST, variance-stabilized transformed counts, approaching a log-transformation).

We next performed mRNA-seq of euploid, T21 and T21-XIST hiPSCs to confirm high uniformity in pluripotency genes over lineage-specific markers [Fig. S2A], and to quantify T21 dosage correction by *XIST* in the presence, absence or upon withdrawal of dox [Fig. 1A,C], which tests whether gene silencing is maintained once established. *XIST* was uniformly expressed in all three T21-XIST clones at 3, 6 and 9 weeks (wks) of dox treatment (“+dox”), and returned to the (“—dox”) baseline upon dox withdrawal [“w/d”, Fig. 1A,C], consistent with *XIST*-negative FISH in w/d samples [Fig. 1B].

To accurately measure chr21 dosage not only by standard differential expression analysis but also chr21 allelic representation, we additionally sequenced euploid and T21 whole genomes with linked-reads to identify and phase variants. In both euploid and T21 hiPSCs, most variants can be grouped in large phase blocks (N50 lengths of 1.8 Mb and 2.9 Mb, respectively), including across chr21, where thousands of heterozygous variants distinguish the *XIST*-bearing (chr21^P^) copy. Allele-specific phasing of mRNA-seq reads indeed reveals that one allele was under-represented with a mean lesser allele frequency (LAF) of one-third for T21 samples. This is in keeping with the presence of a single chr21^P^ copy, and two copies of chr21^M/M2^, whereas the same chr21^P^ allele comprised about half of all allelic reads in the D21 samples [Fig. 2A]. In the absence of dox (-dox), the *XIST*-bearing chr21^P^ copy (T21-XIST) still represented ∼one-third of variant-mapping reads, but was largely depleted in the +dox and w/d samples, which were dox-treated (3wks) and collected after another 3 or 6 wks without dox. These data demonstrate that *XIST* silences most genes on the chr21^P^ copy near-completely, and irreversibly, consistent with our expectations based on mouse *Xist* reports (reviewed in ^22^).

**Figure 2:**
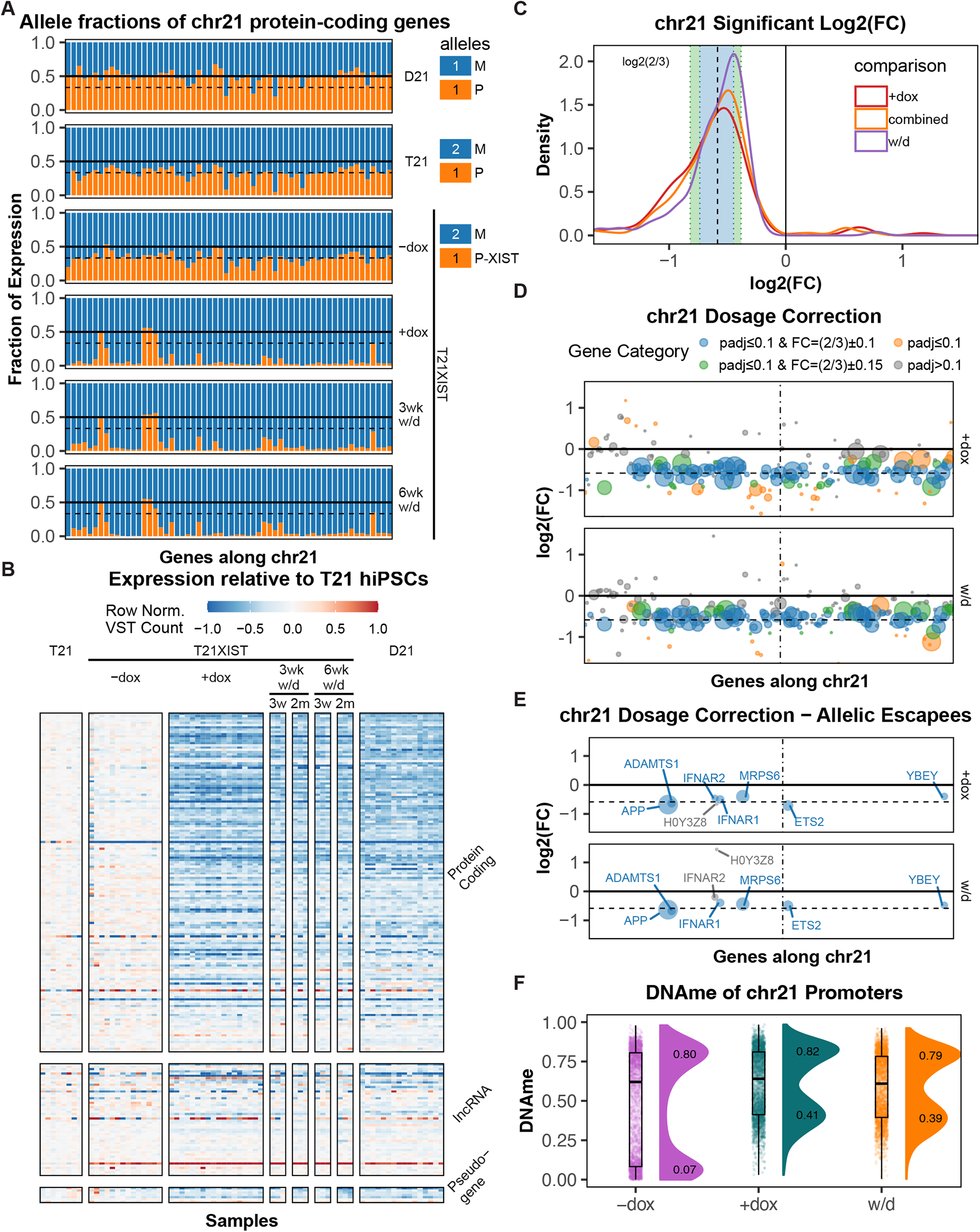
*XIST*-mediated silencing extends across entire chr21P and attracts DNAme. (A) Representation of M/M2 and P alleles of all protein-coding chr21 genes across dox conditions and genotypes. Thick black line at 0.5 marks expected biallelic expression in euploid (D21) cells, and dashed thin black line expected 1/3rd representation for chr21P allele in T21 cells. (B) Heatmap showing VST counts normalized to the mean count in T21 samples (left panels) for all expressed genes on chr21. (C) Distribution of log2(FC) (log2-foldChange) relative to no-dox T21 samples, for all differentially expressed genes (DEGs, p.adj≤0.1). “Combined” represents the differential expression condition used in subsequent figures. Blue and green shaded bands correspond to ±10% and ±0.15 variation from a log2(FC) of -0.59 (dashed vertical line, log2FC expected for complete T21 dosage correction). (D) Log2(FC) values across chr21 genes (arranged by position) compared to T21 samples. Bubble colors shaded as indicated in the legend above, bubble size reflects base mean expression. (E) As in (B), displaying only genes that maintain a LAF ≥ 1/6th in allelic T21 mRNA-seq in +dox or w/d conditions. (F) Distribution of DNAme values from promoter-associated methylEPIC probes on chr21 genes in T21-XIST no-dox, +dox, and w/d. Bimodal distribution medians of probes > and ≤ 60% methylated displayed on their respective peaks.

Likewise, differential expression analysis across chr21 reveals that nearly all genes with assessable expression were depleted by ∼one-third (log2FC of -0.59 ± 0.15), irrespective of their linear distance from the *XIST* transgene, and maintained that expression level even after 3-6 wks of dox withdrawal [Fig. 2B-D]. Altogether, only a very small group of protein-coding genes retained a LAF > 1/6^th^ (*ADAMTS1, APP, H0Y3Z8, IFNAR1/2, MRPS6, ETS2* and *YBEY*), most likely due to mis-phased variants. Indeed, these genes were significantly downregulated by the expected degree (p ≤ 0.1, log2FC -0.59 ± 0.15), except for *IFNAR2* and the adjacent ORF *H0Y3Z8,* which appeared to escape *XIST*-mediated repression by differential expression (p.adj > 0.1, log2FC > -0.3). To compare this efficacy to previously reported T21-*XIST* systems ^21,27^, we considered the degree of silencing across all mutually assessed chr21 genes in prior reports, irrespective of adjusted p-value cutoff, as our study included a greater number of samples. Indeed, the distribution log2FC values in our T21-*XIST* indicates more complete gene repression than in previously published datasets [Fig. S2B].

Comparing differential expression in hiPSCs after 3 wk and 6wk of dox withdrawal, we find that silencing was maintained in w/d samples, irrespective of the length of original dox treatment [Fig. 2B, Fig. S2C]. We therefore assessed DNA methylation (DNAme) at promoters, which reflects stable gene silencing known to remain intact even upon *Xist* deletion in mouse fibroblasts ^45,46^. Excluding likely inactive promoters with high DNAme (>60% methylated), virtually all chr21 promoter-associated CpG probes of the methylEPIC array (Illumina) increased from a median of 7% methylated in the absence of dox (-dox) to 41% methylated after ≥3 wks in +dox [Fig. 2F]. This level is consistent with the presence of a fully methylated alongside two unmethylated alleles (1/3) and is maintained ≥3 wks after dox withdrawal (w/d, 39% methylated), which is also reflected in the absence of reactivated genes by differential and allelic expression [Fig. 2A-E].

We next turned to transcriptome-wide effects of T21 and *XIST*-corrected gene dosage. While levels of chr21 genes clearly increased in T21 relative to D21 and decreased in T21-XIST +dox relative to -dox samples, all other chromosomes also responded to chr21 dosage, but roughly balanced up- and down-regulated genes [Fig. 3A]. Consistently, samples with an effective chr21 dosage matching euploid samples (D21, and +dox or w/d T21-XIST) segregated from T21 and no-dox T21-XIST samples in PCA [Fig. 3B]. Because our experimental design included the original, non-transgenic D21 and T21 hiPSCs (198-1/2/5) with all treatments (±dox and w/d), we were able to observe that their acute +dox treatment triggered expression changes that correlated to some extent with the T21 dosage effect [Fig. 3C left], which was still significant after excluding chr21 genes from the correlation [Fig. S3A, left]. This was not unexpected, because doxycycline can partially impact mitochondrial translation in eukaryotic cells ^47^, and mitochondrial pathways are also altered in DS ^48–50^. We therefore adjusted for this small dox-specific effect in our analysis (see methods). We then compared T21 over D21 log2FC expression changes with the combined (+dox and w/d) *XIST* correction, which revealed a highly significant and negative Pearson correlation of -0.5 [Fig. 3C, right] that persists even when chr21 genes are excluded [Fig. S3A, right]. These data indicate that differentially expressed genes (DEGs) responding to excess chr21 dosage are also changing in the opposite direction when excess chr21 dosage is corrected via *XIST*. This large-scale reversal was also evident in gene-set enrichment analysis (GSEA), with a dominance of anti-correlated terms [Fig. 3D, top left and bottom right quadrant], many of which related to apoptosis, stress, ER, mitochondria and translation terms. Plotting the top three most-concordant terms for each of these categories [Fig. 3E], T21 and dosage-corrected samples were near-uniformly opposed in biological processes previously implicated in DS, including apoptosis ^17,51,52^, oxidative stress via NRF2 ^16,53^, mitochondria ^18,48–50^, and translation ^14,54^. Importantly, serine biosynthesis pathways, recently attributed to a general response to aneuploidy ^20^ were also highly represented, indicating that serine metabolism returns to baseline upon T21 dosage correction, despite the continued presence of the supernumerary chr21 copy.

**Figure 3:**
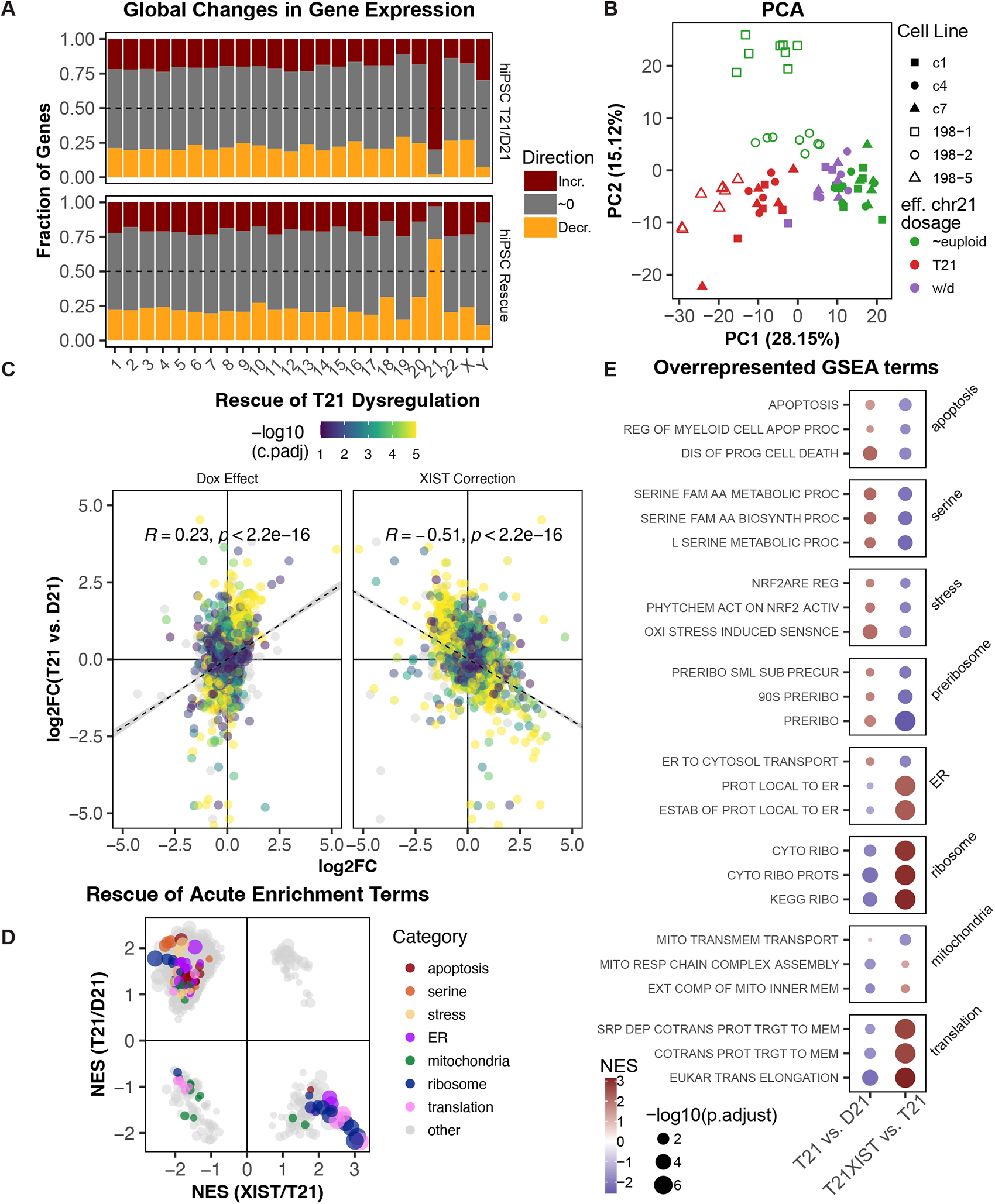
Transcriptome-wide response to T21 gene dosage and correction via *XIST*. (A) Fraction of genes differentially expressed in response to T21 (top, relative to euploid) and T21-XIST dosage correction (bottom, relative to T21 and no-dox) by direction (increasing, decreasing, or unchanged, ∼0) and by chromosome. (B) PCA of all mRNA-seq samples, batch-corrected for timepoint and dox status. Open symbols for original euploid (198-1/2) and T21 (198-5) lines, filled symbols for T21-XIST transgenic lines (c1/4/7). Colors indicate effective chr21 dosage as indicated in Figures 1 and 2 (w/d colored separately to distinguish from +dox T21-XIST samples). (C) Overlap of differential expression between hiPSC T21/D21 (y-axis) over the dox effect (left, non-transgene +dox/no-dox) or T21-XIST dosage correction (right, T21XIST+dox & withdrawal / T21). Adjusted p-values combined by Fisher’s method (C.padj) for all DEGs, and colored by -log10(C.padj) provided C.padj ≤0.1. Linear regression plotted with the Spearman correlation coefficient and p-value for all genes. (D) Overlap of enriched GSEA terms comparing normalized enrichment scores (NES) for hiPSC T21/D21 over dosage-corrected T21-XIST/T21. Bubbles correspond to the -log10(combined pvalue by Fisher’s method). Colors denote chosen categories (all with combined pvalue ≤ 0.2). (E) Individual significantly-enriched terms (combined p≤0.1) from each of the categories in (D), selected by concordance measure (Euclidean distance from origin in D). Colors denote NES, bubble size the individual -log10(p.adj).

### T21 gene dosage effects in mature neural cell types

To assess T21 gene dosage effects in neural lineages, we applied a well-established monolayer differentiation protocol that mimics *in vivo* neural development in the establishment of neurogenic niches (rosettes) and in the production of both mature neurons and astrocytes ^55,56^. We chose this approach as nearly all T21 hiPSC-derived neuronal RNA-seq data reported to-date was generated from immature neurons in the absence of astrocytes. We performed mRNA-seq in 10-12 wk neural cultures, obtained 5-6 wks after low-density re-plating of 4 wk NPCs [Fig. S3B]. T21 neural cultures showed elevated chr21 transcript levels with a peak near the expected +0.59 log2FC relative to euploid samples [Fig. 4A,B]. As expected, neural and hiPSC DEGs and GSEA-enriched terms were positively correlated, while neural GSEA terms were largely anti-correlated in direction with terms enriched in XIST-corrected hiPSCs [Fig. 4C-E], which also extended to the three most-concordant terms across all comparisons [Fig. S3B]. Overall, we find *XIST-*mediated T21 dosage correction is reflected in a majority of significantly enriched terms differentiating T21 hiPSCs and neural cultures from their euploid controls [Fig. S4].

**Figure 4:**
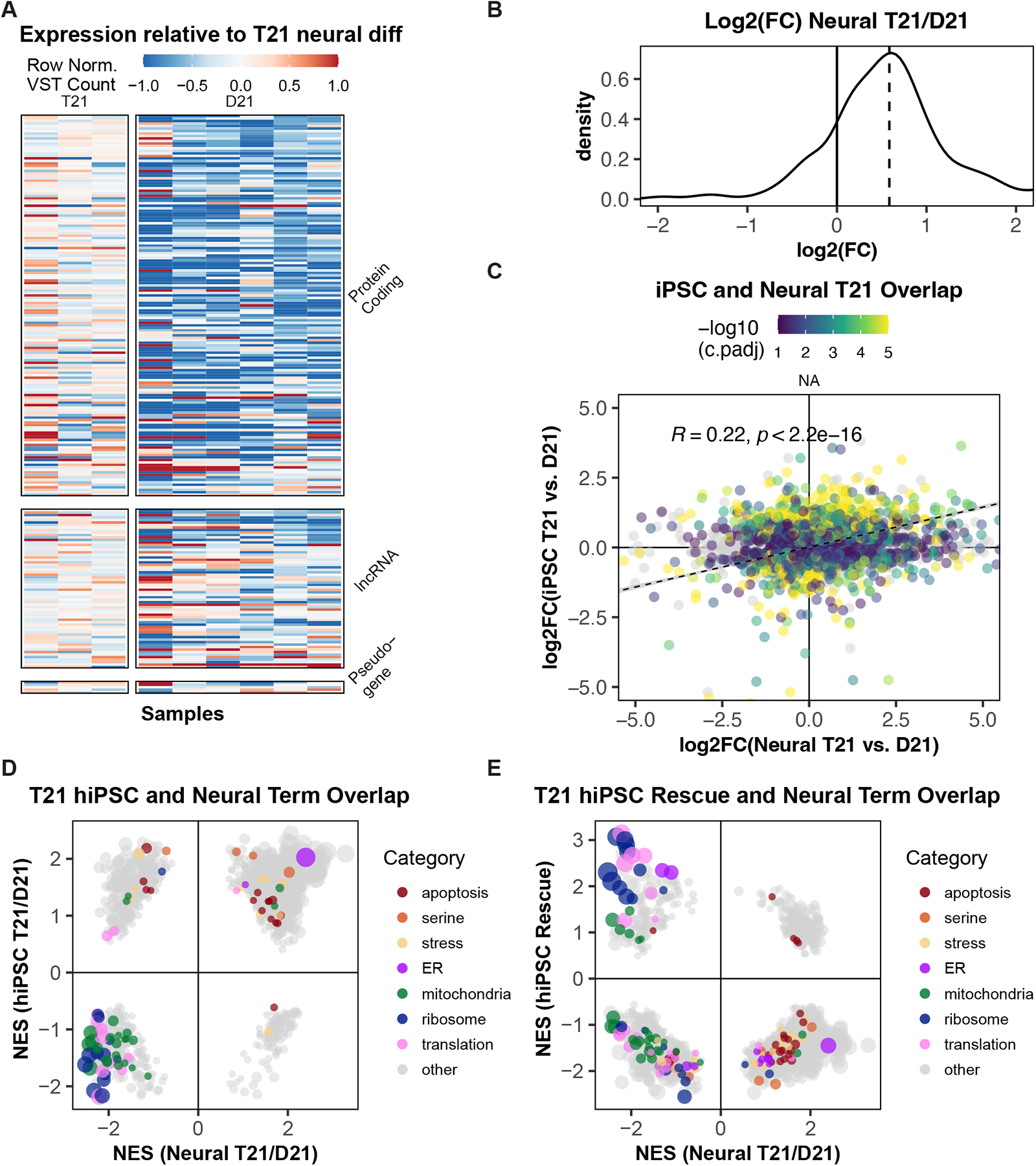
Concordant T21 dosage imbalance and transcriptomic response after neural differentiation. (A) Heatmap showing VST counts normalized to the mean count in T21 neural samples (left panels) for all expressed genes on chr21. (B) Distribution of log2(fold-change) from chr21 genes comparing neural T21 to D21. Dashed black line denotes expected log2(FC) of +0.59 (includes all chr21 genes, irrespective of p.adj). (C) Overlap of differential expression between hiPSC T21/D21 (y-axis) over neural T21/D21 samples (x-axis). Adjusted p-values were combined and colored as in Fig. 3C, with linear regression line (dashed, with Spearman coefficient and p-value for all genes with c.padj ≤ 0.1). (D) Overlap of enriched GSEA terms comparing normalized enrichment scores (NES) for hiPSC T21/D21 (y-axis) over neural T21/D21. Bubble plots corresponds to the -log10(combined pvalue by Fisher’s method). Colors denote chosen categories (all with combined pvalue ≤ 0.2). (E) As in (D) but combining neural T21/D21 GSEA (x-axis) with GSEA of dosage-corrected T21-XIST hiPSCs (hiPSC rescue) on the y-axis.

To assess whether the number of over-represented GSEA terms relating to cell death and apoptosis in T21 neural cultures manifested in differential neuronal survival, we employed a recently developed red genetically-encoded cell death indicator (RGEDI) and tracked two neuronal subtypes by automated longitudinal imaging [Fig. 5A, Fig. S5A-D] ^57,58^. Both T21 interneurons (hazard ratio of 4.08) and T21 forebrain neurons (hazard ratio of 2.57) demonstrated significantly increased cell death (RGEDI/EGFP ratio > 0.15, Holm-corrected p<0.01, linear mixed model) over ∼10 days of single-cell tracking, compared to euploid controls [Fig. 5B,C]. Additionally, we also assessed live neurons (RGEDI/EGFP ratio < 0.15) and noted that there was a significant (p≤10^-8^) reduction in EGFP signal in T21 compared to euploid controls [Fig. S5E], that mirrored the down-regulation of ribosome and translation-related GSEA terms in both T21 hiPSCs and neural cultures [Fig. 4D, Fig. S4].

**Figure 5:**
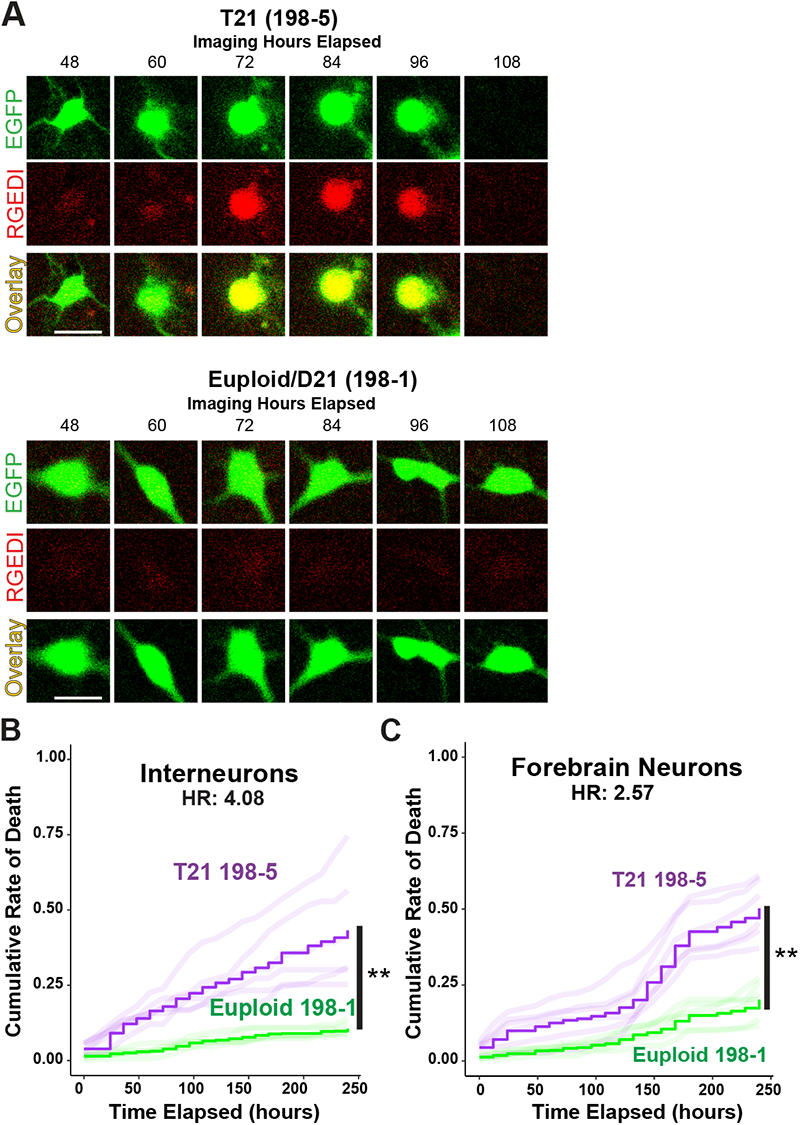
Increased cell death of T21 relative to euploid interneurons and forebrain neurons. (A) Representative sequence of a TS21 interneuron labeled with RGEDI-P2a-EGFP dying over the course of imaging and a control neuron surviving during the course of imaging (scale/width is 10 μm). (B) Plot of the cumulative rate of death of TS21 and control interneurons. Step functions correspond to the mean cumulative death rate across wells per line and the faint lines represent each well replicate (n = 6 wells each, HR = 4.08, Linear mixed model with Holm correction, ** *p* <0.01). C) as in B) for the cumulative rate of death of TS21 and control forebrain neurons (n = 6 wells each, HR = 2.57, Linear mixed model with Holm correction, ** *p* <0.01).

### T21 dosage correction early in neural differentiation rescues neurogenesis

Staining for neuronal MAP2 and astroglial S100B markers, we observed an excess of astrocytes in the T21 relative to euploid neural differentiations. Both neurons and astrocytes derive from NPCs ^59^, and astrogliosis has been reported in DS models ^33,34^ and fetal samples ^28,30,32^, where T21 NPCs prematurely shift from neurogenesis to astrogenesis. To determine whether *XIST*-mediated T21-dosage correction could prevent this lineage skew, we first confirmed that *XIST* expression and H2AK119ub enrichment would persist in 4 wk NPCs, which was the case for well over 75% of NPCs derived across three independent clones, treated with +dox as hiPSCs [Fig. 6A]. Indeed, even NPCs derived from untreated T21-*XIST* hiPSCs but receiving dox from the first day of differentiation (“d0”) presented with *XIST* and H2AK119ub foci in ≥75% and ≥65% of all cells, respectively. Co-staining for MAP2 and S100B on day28 (4 wk) of neural differentiation revealed a consistent and highly reproducible astrogenic skew in T21 and no-dox T21-XIST samples relative to isogenic euploid control lines across multiple rounds of differentiation [Fig. 6B]. Remarkably, neurogenesis was fully rescued in neural differentiations from pre-treated as well as d0+dox treated hiPSCs, across all three independent T21-XIST clones. Because the overall sum of committed cells expressing either marker did not differ significantly between T21 and euploid or T21-XIST dosage-corrected samples, we propose that T21 dosage does not suppress NPC commitment in general (as suggested ^27^), but rather advances the shift to astrogenesis, which is consistent with reports in fetal DS and rodent DS models ^30,34^.

**Figure 6:**
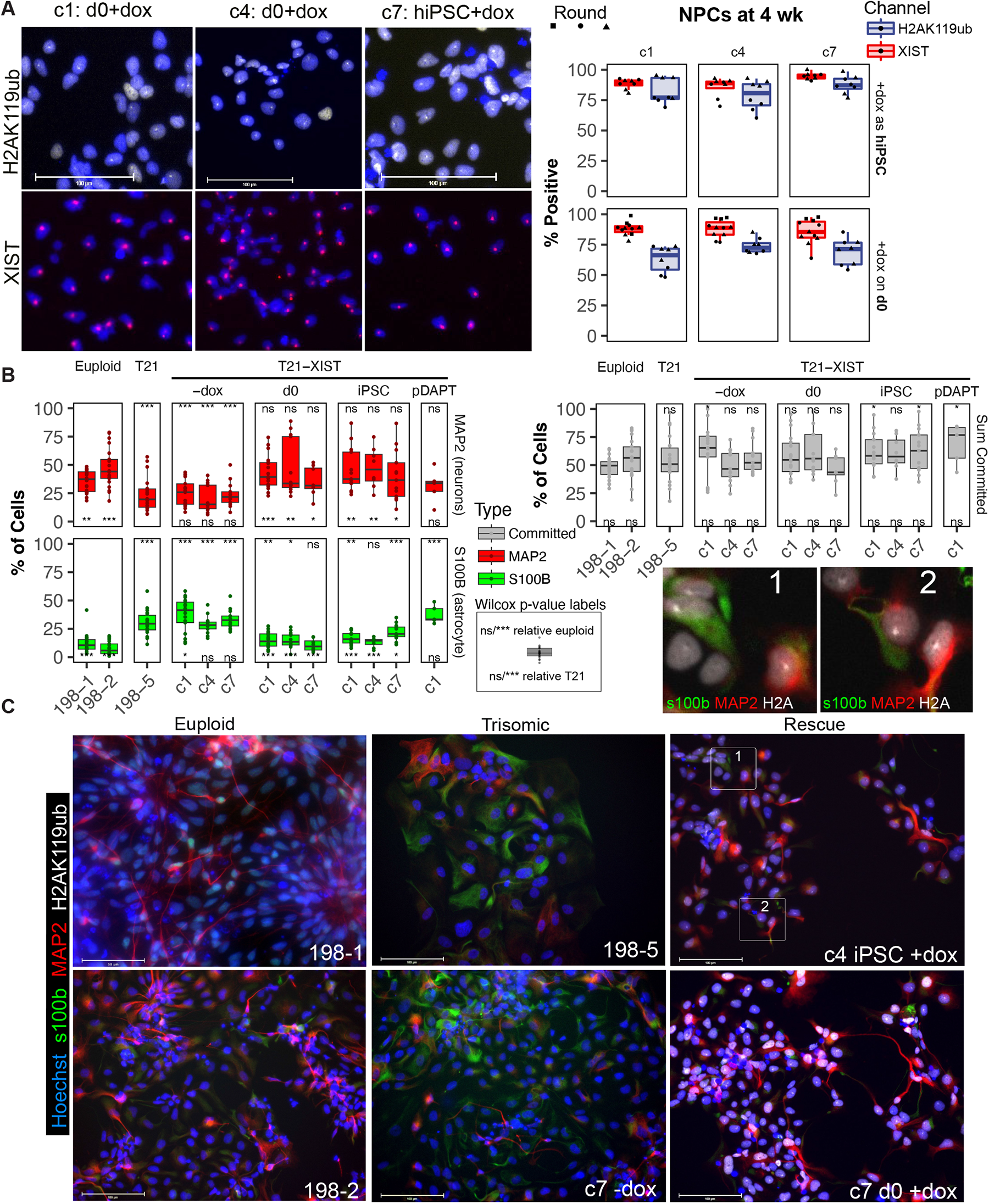
Early astrogenesis at neurogenic expense is suppressed by T21 dosage correction prior to NPC stage. (A) Representative images (left) and quantification (right) of H2AK119ub (IF) and XIST (FISH) signal after neural differentiation from dox treated cells. Dox treatment initiated while cells were hiPSCs (left top), or on the first day of differentiation (“d0”, left bottom). (B) Relative lineage commitment to astrocytes (S100b) or neurons (MAP2) in euploid, T21, and T21-XIST cells under dox treatment. Symbols (Wilcoxon two-tailed p-value level, or “ns” for p > 0.1) above each boxplot denote significance of median difference from the corresponding euploid (198-1/2) condition, whereas symbols below each boxplot compare to T21 (198-5) lineage proportions. (C) Representative images at d28 of neural differentiation in euploid (left column), trisomic (center column) and rescued (right column) labeled for neuronal MAP2 (red), astroglial S100B (green) and H2AK119ub accumulating on chr21 (white). Insets (labeled 1,2) display H2AK119ub foci in both astrocytes and neurons.

To test whether this interpretation is replicated *in vivo,* at the very onset of cortical astrogenesis, we turned to the Ts65Dn model [Fig. S6]. At embryonic day 18.5 (E18.5), the density of S100B+ positive astrocytes in the developing cortex was significantly elevated in Ts65Dn embryos over their wildtype litter mates (p < 0.05). In contrast, there was no significant difference in the density of cycling Ki67^+^ cells in the proliferative zone [Fig. S6], indicating the early cortical astrogenesis in Ts65Dn embryos was not associated with delayed commitment and continued proliferation of neural progenitors, but rather an inherent skew towards astrogenesis also observed in other brain regions ^30,34^.

### Postmitotic neural cell types still induce *XIST* and attract repressive chromatin marks

Given the high rate of *XIST* induction in T21-XIST NPCs that only received dox at the onset of differentiation (d0), we next explored whether the *XIST* transgene would remain dox-responsive in terminally differentiated cells [Fig. 7A]. To this end, we forced NPCs (∼4 wk neural differentiation) out of the cell cycle into terminal differentiation by inhibiting the Notch signaling pathway for 5 days with gamma-secretase inhibitor DAPT, and added dox a day after completing DAPT treatment (d33). To confirm DAPT treatment was sufficient to deplete mitotic cells, we also labeled cells transiting S-phase with nucleotide analog EdU, and co-stained for this label and *XIST* FISH after two days in +dox media (d35). DAPT-treated NPCs incorporating EdU comprised ≤6% of the neural population on average, which were rarely EdU/*XIST* double-positive (≤26%), whereas *XIST* was induced in ≥40% of all EdU^−^ cells (94% of all cells). We conclude that *XIST* was reliably induced in a sizeable fraction of post-mitotic T21-XIST cells that had terminally differentiated without dox. We next assessed these post-mitotic cells after 3 to 6 weeks of dox treatment post-DAPT exposure (“pD”) to provide sufficient time for H2AK119ub enrichment. We included “d0” controls that were treated with the standard 1 μM dox concentration throughout neural diff, as well as low-dose dox “LD” samples, which were treated with 0.01 μM dox up to the DAPT exposure, and standard 1 μM dox thereafter, to test whether “pre-marking” the TRE3G promoter without effective induction through differentiation may render it more responsive in terminal cell types. On average, over half of d0 and LD cells featured prominent XIST clouds, followed by ≥ 40% of pD samples, across both c1 and c7 clones and in three independent rounds of differentiation. As before, H2AK119ub signal lagged slightly behind *XIST*, but consistently marked 27% – 36% of all cells (58-65% of *XIST+* cells), including MAP2+ neurons and S100B+ astrocytes [Fig. 7C,D].

**Figure 7:**
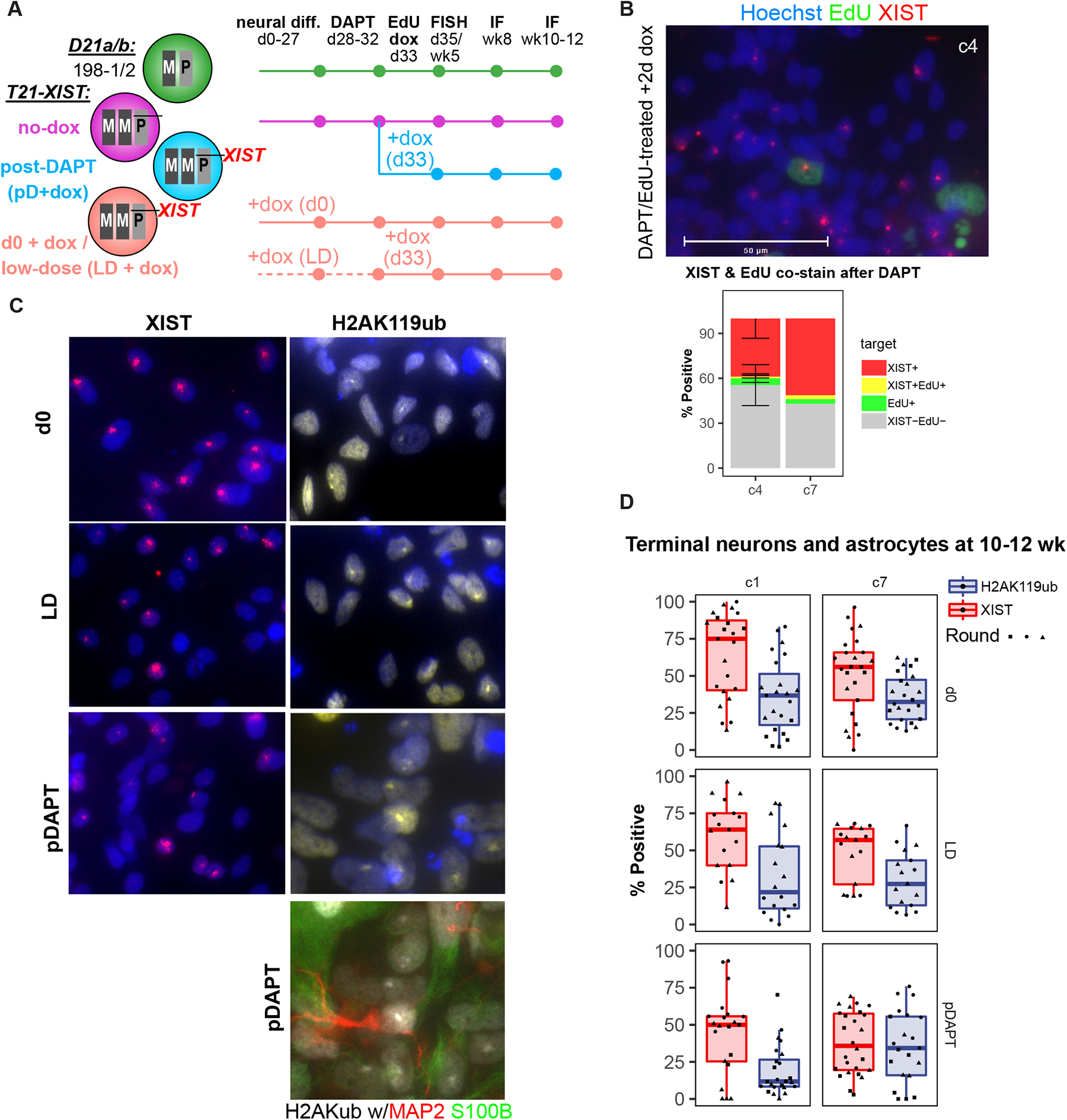
Post-mitotic neurons and astrocytes can induce *XIST* and deposit H2AK119ub after terminal lineage commitment. (A) Sample and timeline schematic of XIST induction during (d0+dox) and after terminal neural differentiation post DAPT treatment (pD+dox). Cell cycle exit of NPCs is assessed by EdU-labeling of S-phase transiting cells, with co-detection of *XIST* by FISH after 2 days in dox media. Chr21 Barr body formation (H2AK119Ub) and lineage commitment (MAP2/S100B) are assessed by IF in 8 wk and 10-12 wk cultures. (B) Representative *XIST* (red) and EdU (green) image and quantification of d36 neural differentiation following 4d of DAPT and 2d of dox treatment. Less than 6% of cells were EdU+ (green), EdU+XIST+ cells represented less than 2% of total cells. (C,D) Representative images (C) and quantification (D) of H2AK119ub (yellow) IF and XIST-FISH (red) signal after neural differentiation from different dox treatments. Bottom pDAPT panel shows H2AK119ub IF (white) co-stained with MAP2 (red) and S100B (green).

### *XIST* induction in terminal neurons and astrocytes corrects T21 gene dosage

We next performed single-nuclei RNA-seq, to determine whether such *XIST* induction and H2AK119ub accumulation in these terminally differentiated neural cell types reflect T21 dosage correction [Fig. 8]. Transgenic neural differentiations treated with dox from d0 or pD were collected alongside untreated T21-XIST c7 cells (“T21”), and two euploid lines (198-1/2 or “D21a/b”). Nuclei distributed across four broad clusters, of which three corresponded to the expected NPC, neuronal and astroglial populations [Fig. 8A, S7A]. Cell cycle analysis classified the smaller NPC cluster as S and G2/M phase cells, while all other clusters scored as G1, consistent with terminally differentiated G0 cells [Fig. 8A, S7A]. We labeled the first three clusters according to their predominant cell type using cell type markers from ^60^, and the fourth as “Other” given lower scores returned by sctype ^61^. Due to its proximity to higher-scoring oligodendroglial progenitor cells (OPC), this fourth cluster probably reflects OPC-derived lineages that predominantly arose from one of the euploid lines (D21a). Crucially, *XIST* induction in T21-XIST neural differentiations (d0 & pD) was detected in all clusters with 74-94% of d0+dox treated and 24-58% pD+dox treated cells [Fig. S7A], in agreement with our FISH and IF data [Fig. 7].

**Figure 8:**
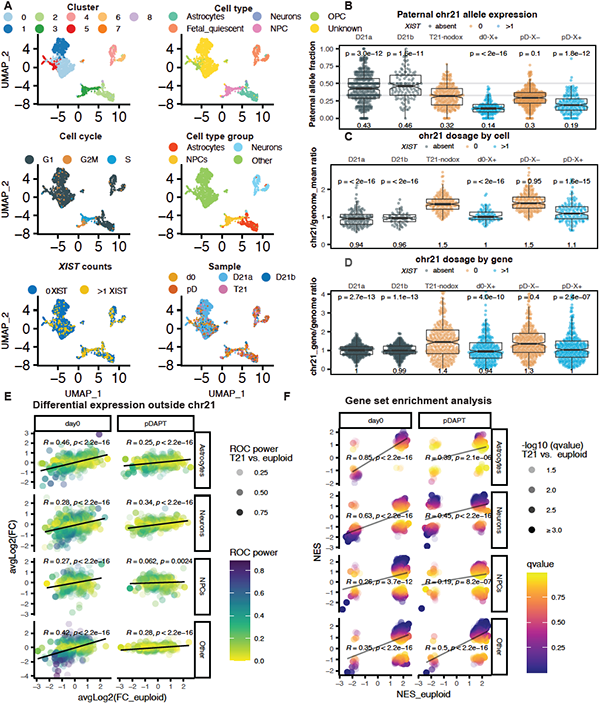
Effective monoallelic repression of chr21 genes by *XIST* induced in terminal neurons and astrocytes. (A) Single-nuclei RNA-seq UMAP of terminal 10-12 wk neural cultures (as outlined in Fig. 7A). Integrated UMAP colored by original clusters, granular and grouped cell types, cell cycle stage, XIST count categories and samples. (B) Paternal allele fraction by cell, aggregated over all chr21 genes. Cells are split by sample (D21a/b, T21XIST-nodox, d0+dox, pD+dox) and *XIST* status (0, >1 counts denoted “X-“ and “X+”), except for euploid lines (absent XIST transgene, grey). Median paternal allele fraction and significance of difference in medians relative to T21-nodox cells (Wilcoxon p-value) are denoted below and above boxplot, respectively. (C) Normalized gene expression ratio (chr21 over genome mean excluding chr21) by cell, aggregated over all chr21 genes. Cells are split as in B (excluding cell type group “Other”), with median expression ratio and difference in medians relative to T21-nodox cells (Wilcoxon p-value) denoted above and below boxplot, respectively. (D) Normalized expression ratio by chr21 gene, aggregated over cells. Mean ratios per gene calculated from cells are split as in C. Respective median expression ratios and difference in medians relative to T21-nodox cells (Wilcoxon p-value) denoted above and below boxplots. (E) Differential expression of non-chr21 genes relative to XIST-negative postDAPT and T21-nodox cells. Sample-averaged log2(FC) values in euploid cells (x-axis) correlate with XIST-positive d0 and pDAPT log2(FC) values (y-axis), faceted by cell type group. Colors and transparency denote respectively, ROC power and ROC power of the euploid relative to T21 comparison. Regression line (black), Pearson coefficient and p-value (Fisher-transformed) denoted on each plot. (F) GSEA results of differential expression comparisons depicted in (E), correlating normalized enrichment scores (NES), and plotted as in (E). Plotted MSigDB gene sets limited to Hallmark and Canonical pathways subsets significant (q-value ≤ 0.1) in T21/euploid comparison (x-axis). Colors and transparency denote respectively, -log10-transformed q-values of each comparison and the euploid to T21 comparison. Regression line (grey), Pearson coefficient and p-value (Fisher-transformed) denoted on each plot.

To determine whether *XIST* induction repressed T21 gene dosage, we assessed three complementary metrics. First, we phased single-nuclei RNA-seq reads and summed allele-specific counts across all chr21 genes to calculate the cumulative paternal allele representation, thereby mitigating the effect of transcriptional bursts in individual nuclei [Fig. 8B]. As expected, euploid cells have a significantly higher chr21^P^ allele fraction compared to T21 cells (0.46 vs. 0.32, Wilcoxon p = 1.6×10^-11^), whereas T21-XIST day0+dox treated cells have a significantly lower chr21^P^ allele fraction (0.14, Wilcoxon p < 2.2×10^-16^). Crucially, while *XIST*-negative pD (pD-X^−^) cells show little reduction in this *XIST*-carrying paternal allele, *XIST*-positive pD cells (pD-X^+^) reveal significant repression of this allele (0.19, Wilcoxon p-value = 1.3×10^-12^). Separating cells by cell type group [Fig. S7B] reveals *XIST+* d0 and pD neurons repress the transgene-bearing chr21^P^ copy most robustly, with paternal allele fractions of 0.15 and 0.17 (Wilcoxon p<10^-4^), respectively.

Second, we compared the normalized expression for each gene on chr21 to the genome-wide mean without chr21. We then determined the per-cell cumulative distribution of all chr21 genes, as well as the per-gene cumulative distribution by cell type group [Fig. S7C, D], and aggregated these ratios across all neurons, astrocytes, and NPCs [Fig. 8C,D]. Indeed, euploid and T21-nodox medians fell near the expected 1.0x and 1.5x ratios relative to the genome-wide mean, indicating chr21 genes are not inherently dosage-compensated. Likewise, *XIST*-negative pD nuclei failed to differ significantly from the T21-nodox sample (1.5x and 1.3x for the per-cell and per-gene ratio, respectively). *XIST*+ pD nuclei however displayed near-euploid chr21 dosage medians (ratios of 1.1x and 1.0x, Wilcoxon p-values ≤ 2.4×10^-7^), very close to d0 nuclei [Fig. 8C,D]. Separated by cell type group, neurons again demonstrated near-complete dosage compensation in *XIST+* pD nuclei with median ratios of 0.97x and 0.92x on a per-cell and per-gene basis, respectively [Fig. 8C,D]. Assessing both metrics across full distributions for each sample, the *XIST*-negative pD distribution tracked closely with T21-nodox cells, remaining well within its 95% confidence interval. In contrast, the *XIST+* pD and d0 distributions shifted to the left significantly (dashed lines, Kolmogorov-Smirnov, K.S. test p-values) towards the euploid distributions for all cell type groups, irrespective whether normalized expression ratios were averaged over cells or genes [Fig. S7C, D]. Aggregating these distributions across astrocytes, neurons and NPCs illustrates *XIST+* d0 and pD nuclei tracked on top of each other, with their 50% percentile ratios near-identical to euploid nuclei [Fig. S7E,F]. In sum, these data indicate that *XIST* corrects T21 gene dosage in post-mitotic cell types via robust repression its chr21^P^ copy, especially in neurons, whereas NPCs mitigate *XIST’s* impact to some extent, likely due to continued passage through M phase.

As our third metric for T21 dosage correction, we performed differential expression analysis on variance-stabilized data ^62^, and compared effectively trisomic (*XIST*-negative pD and nodox) nuclei to euploid (D21a/b), *XIST+* pD and d0 nuclei for each cell type. Indeed, generally lower expression of chr21 in euploid relative to trisomic nuclei was also observed for *XIST+* pD and d0 nuclei (Fig. S7G). Due to greater dispersion inherent in single-nuclei RNA-seq, the moderated effect sizes (avg.log2FC) reflected in variance-stabilized data were smaller than in our bulk RNA-seq (Fig. 2,4) but enabled comparing euploid to *XIST*-corrected T21 gene dosage. Interestingly, *XIST* repressed chr21 genes to the near-euploid median, in all except ‘Other’ cell types, which showed a small but significant residual excess expression (Wilcoxon p = 0.0007, Fig. S8A). While the linear distance to the *XIST* transgene integration site did not correlate with mean log2(FC) values in any of the d0+dox cell types, there was a distance effect in pD+dox neurons and NPCs [Fig. S8B,C].

Importantly, differential expression of non-chr21 genes in *XIST^+^* d0 and pD nuclei correlated strongly with log2(FC) values in euploid relative to effectively trisomic nuclei across all cell type groups [Fig. 8E], with Pearson coefficients ranging from 0.25-0.46 in all except *XIST^+^* pD NPCs. GSEA also reflected such transcriptome-wide reversion with strong NES correlations between *XIST+* d0, pD and euploid conditions in MsigDB Hallmark and canonical pathways that were significantly enriched (q-value ≤ 0.1) in euploid relative to effectively trisomic nuclei [Fig. 8F]. Again, the correlation in NPCs was lowest across all cell type groups, yet still significant (R=0.19, p=8.2×10^-7^). Importantly, the vast majority of MsigDB Hallmark and canonical pathways significantly enriched in any pairwise comparison (combined Fisher’s method p≤1×10^-4^) followed the direction of change in the remaining third comparison [Fig. S8D]. In sum, these results indicate that the degree of chr21 dosage correction in *XIST+* d0 and pD nuclei is sufficient to revert T21 cells towards the euploid expression profile. We conclude that this new T21-*XIST* transgene corrects chr21 dosage even in terminal and post-mitotic cell types, which will facilitate detailed exploration of cellular and transcriptional T21 hallmarks that remain reversible after completed development.

## DISCUSSION

The relevance of the inducible T21-XIST system presented herein ranges from DS biology and cellular responses to aneuploidy in general, to fundamental questions relating to human *XIST*-mediated gene silencing and escape. In hiPSCs, our T21-XIST system is induced with near-uniformity [Fig. 1] and corrects T21 dosage to euploid levels across chr21 [Fig. 2], which we demonstrate by inclusion of isogenic euploid control hiPSCs lines we derived in parallel from a male donor with mosaic T21 ^36^. Leveraging phased sequence variants from linked-read WGS, we can assign mRNA-seq reads allele-specifically to show that chr21 genes derive half of their expression from the chr21^P^ allele in euploid (D21) cells, and one-third in T21 cells, but nearly extinguish expression from this *XIST*-bearing copy after dox induction [Fig. 2].

*XIST*-mediated T21 dosage correction extends to genes responding to chr21 dosage on all chromosomes, reflecting a transcriptome-wide reversion of gene expression that moves dox-treated T21-XIST hiPSCs towards their isogenic euploid controls [Fig. 3]. A balanced experimental design and stability of T21 silencing after dox withdrawal (w/d) enabled us to adjust for dox effects that can inhibit mitochondrial translation ^47^ and selenocysteine incorporation ^63^. GSEA reflects the large-scale reversion of T21-responsive cellular components, pathways, and functions [Fig. 3], including boosted expression of serine biosynthesis genes. These genes are notable as dermal fibroblast of DS, as well as Patau (T13) and Edwards (T18) syndromes, depend on serine for proliferation ^20^. As such genes return to baseline upon correction of T21 dosage [Fig. 3], despite maintenance of the chr21 copy number, metabolic demand for serine, while common to all three viable human autosomal aneuploidies, results from expression of the supernumerary chromosome rather than its presence. We also find resurgent expression of ribosomal protein genes and translation factors in T21-dosage corrected hiPSCs [Fig. 3], which may be due to a resetting of the integrating stress response ^14^. Indeed, cellular stress pathways and apoptosis are dampened in response to T21 dosage correction, which is consistent with the previously reported excess of apoptosis ^17,51,52^, mitochondrial dysfunction ^18,48–50^ and oxidative stress ^16,53^ in DS.

In T21-driven expression of neural cultures, we find many of the same terms are enriched by GSEA and concordant with T21 hiPSCs [Fig. 4]. In keeping with a recent DS meta-analysis ^64^, the interferon response [Fig. S4], as well as mitochondrial pathways and apoptotic terms [Fig. S3], are particularly exacerbated in T21 neural cultures. Consistent with these results, we demonstrate impaired survival of T21 forebrain neurons and interneurons relative to isogenic euploid controls [Fig. 5], as well as reduced translation of EGFP [Fig. S5]. Because GSEA-enriched terms and related genes responded to *XIST*-mediated T21 dosage correction in hiPSCs, they likely reflect cellular phenotypes that depend on ongoing excess chr21 expression, and may therefore remain amenable to T21 dosage correction after completed development.

During early neural differentiation, we observe that T21 NPCs produce an excess of astrocytes at the expense of neurons [Fig. 6], which is consistent with astrogliosis noted in fetal DS ^28,30,32^. Because *XIST*-mediated T21 silencing remains inducible during differentiation and persists into the NPC stage, neurogenesis is fully rescued when T21-dosage is corrected [Fig. 6]. This is consistent with a prior report using T21-*XIST* dosage correction ^27^, but we can attribute this neurogenic rescue to suppression of a conserved T21 astrogenic bias, rather than slowed commitment of T21 NPCs [Fig. 6]. We replicate this observation at the onset of cortical astrogenesis in Ts65Dn embryos (E18.5), which indeed show a significant increase in S100B+ cells relative to wildtype embryos, while the density of Ki67+ neural progenitors remains unchanged [Fig. S6]. Several chr21 genes have been suggested to play a role in this astrogenic lineage bias, including amyloid precursor *APP, S100B*, and *DYRK1A* ^30,34^. In addition, it is possible that cellular stress pathways impact NPC lineage decisions, as seen in rodent models of neurogenesis ^65^. One possible culprit may be oxidative stress that is elevated in DS hiPSC-derived neurons and NPCs ^40,66^ and also implicates these and additional chr21 genes ^16,30,53^.

We also examined T21-*XIST* hiPSC-derived mature neural lineages after forcing NPCs to terminally differentiate and thus commit to terminal neural cell fates via notch-inhibitor DAPT [Fig. 7A,B]. This more mature (∼9-12wk), post-mitotic neural population is composed of terminally differentiated neurons and astrocytes, as well as residual neural progenitors and some oligodendroglial lineages [Fig. 8, S6]. Remarkably, a large fraction of this terminal post-mitotic population can induce *XIST* and attract repressive H2AK119ub on chr21 in response to dox [Fig. 7], regardless of whether they were never exposed to dox prior to cessation of the DAPT treatment (post-DAPT, “pDAPT/pD”), or were treated with only a low (“LD”, 1% of the standard 1 μM) dox concentration prior to DAPT treatment. In contrast, a previously reported T21-*XIST* system could not be induced at all anymore in post-mitotic monolayer neural cultures ^27^.

Our results indicate our T21-*XIST* system represents a major improvement, which we attribute to technical characteristics of transgene expression. First, our system uses a *UBC-rTTA* construct integrated at the AAVS “safe-harbor” locus, in contrast to prior studies that used EF1a ^21^ or CAG promoters ^27^ at this locus or re-delivered a high-copy *EF1a-rTTA* construct via piggyBac to NPCs ^67^. Second, there may be sequences differences in these human *XIST* cDNA constructs that could impact the mature *XIST* RNA. And third, we integrated our *XIST* cDNA transgene outside of *DYRK1A*, to minimize its impact on *DYRK1A* prior to dox treatment, but also to avoid attracting gene-body associated DNAme from overlapping transcription, which serves to suppress intragenic transcription start sites ^68^.

We assessed whether *XIST* induction in terminal, post-mitotic neural cell types (“pD”) was as effective in correcting T21 dosage as it was during differentiation (“d0”) using three independent analytic approaches: chr21^P^ allele representation, normalized expression across chr21, and differential expression across the transcriptome [Fig. 8, S6, S7]. The chr21^P^ allele is effectively repressed to nearly the same degree in *XIST+* pD as in d0 cells, especially in neurons. [Fig. 8B,S6]. Likewise, both the median sum of normalized chr21 expression by cell, as well as the entire distribution of cells, shift toward the euploid mean ratio [Fig. 8, S6]. The same significant shift is observed when aggregating the mean ratio across cells for individual genes [Fig. 8, S6]. While some residual cycling NPCs retain excess expression in a subset of chr21 genes far (∼20 Mbp) from the *XIST* integration site [Fig. S7], both normalized expression and allele-specific metrics indicate *XIST* represses T21 dosage across all cell type groups, being most effective in post-mitotic neurons. The transcriptome-wide analysis mirrors these results: differential expression in *XIST^+^* d0 and pD cells correlates significantly with that of euploid cells when compared to non-induced, *XIST*-negative T21 cells [Fig. 8E]. Likewise, canonical pathways significantly enriched by GSEA in *XIST^+^* d0 and pD cells match and follow the direction of change observed in the euploid to T21 comparison [Fig. 8, S7]. In sum, these data demonstrate for the first time that post-mitotic terminal neural cell types remain responsive to chr21 gene dosage. This motivates several avenues for future investigation, for example to dissect neuron-intrinsic – and extrinsic phenotypes ^69^ that will be relevant for developing cell-type specific interventions ^70^, or to identify pathways that remain amenable to novel therapeutic approaches ^14^.

Regarding *XIST* biology, our results suggest human neural lineages remain competent to initiate *XIST*-mediated heterochromatin deposition even non-cycling cells, provided *XIST* can be induced. Similar developmental plasticity in initiating ^25,26^, or re-initiating ^71,72^ *XIST*-mediated repression was previously reported only for proliferative hematopoietic cell lineages. Our work therefore suggests *XIST* may be able to re-establish X chromosome inactivation even outside pluripotency and hematopoiesis, which is relevant to recently reported X chromosome conformational changes in the female brain across the estrous cycle ^73^.

Additionally, prior reports of *XIST* induction in human HT1080 fibrosarcoma cells indicate that spreading of autosomal gene silencing remains reversible ^74^ and remains only effective near its site of integration ^75^. Both of these observations may be impacted in part by *XIST* repressing haploinsufficient genes, which are not uncommon on mammalian autosomes, making whole or partial trisomy a valuable research tool to better understands the determinants of *XIST* spreading ^76^. In hiPSCs, our T21-XIST system repressed virtually every chr21-linked gene without measurable impact of its linear distance from the *XIST* integration site across a large span of expression values, and in irreversible fashion [Fig. 2]. Interestingly, the human inactive X in pluripotent stem cells frequently undergoes progressive reactivation following loss of *XIST* expression ^77^, the order of which we recently showed was determined in part by the distance to the nearest escapee gene ^78^. While our T21-*XIST* dox withdrawal was limited to 3-6 weeks, we found little evidence of reactivation over this period, indicating that DNA methylation served to maintain T21 silencing [Fig. 2]. Whether this remarkable epigenetic stability was due to the paucity of chr21 genes escaping *XIST* to seed potential reactivation, or the levels of *XIST* prior to dox withdrawal, and to which degree these two aspects are related, represent additional areas for future investigation.

## SUPPLEMENTARY FIGURE LEGENDS

**Figure S1**

(A) Taqman copy number (CN) assay for 21q22 (*DYRK1A*) in GM00260-derived hiPSC clones.

(B) Taqman CN assay for 17q25 (*SEC14L1*) in GM00260-derived hiPSC clones.

(C) DNA SureFISH for chr21q, and FastFISH chrX, chr8, chr12, and chr17 centromeres in 198-6/8 with counts and images denoting the dominant staining pattern.

(D) Summary of CytoSNP850k calls across GM00260 and its reprogrammed hiPSC clones. Calls marked with a red ‘x’ were common across all lines, including donor fibroblasts. Call marked with a small ‘x’ were line-specific, except for T21 (denoted by a capital ‘X’).

(E) Absolute difference in allele (theta) frequency of CytoSNP850k probes comparing T21 and euploid clones (D21), colored by T21-specific base calls (red/blue) and hetero/homozygosity (grey and yellow, respectively).

(F) DNA FISH against DSCR8 (Vysis, LSI21) in interphase and metaphase spreads of validated hiPSC clones, with counts and images denoting the dominant staining pattern.

(G) Taqman CN assay for 21q22 (*DYRK1A*) across all AAVS-rTTA subclones of T21 198-5 hiPSCs.

**Figure S2**

(A) Germ layer-specific and pluripotency marker expression across the euploid (198-1/2), T21 (198-5), and transgenic T21-XIST (c1/4/7) hiPSC lines.

(B) Change in log2-transformed expression relative to corresponding T21 samples in this and the two referenced T21-XIST studies. Mean difference of chr21 gene expression comparing w/d T21-XIST samples and T21 samples. Line type denotes the length of the initial dox treatment, and color the duration of dox w/d.

(C) Mean difference of chr21 gene expression between w/d T21-XIST and T21 samples. Line type denotes the length of the initial dox treatment, and color the duration of dox w/d.

**Figure S3**

(A) Same as Fig. 3C but excluding chr21 genes: overlap of differential expression between hiPSC T21/D21 (y-axis) over the dox effect (top, non-transgene +dox/no-dox) or T21-XIST dosage correction (T21XIST+dox & withdrawal / T21). Adjusted p-values were combined (C.padj, calculated using Fisher’s method), and colored by -log10(C.padj) provided C.padj ≤0.1 (or grey for C.padj > 0.1). Linear regression is plotted along with the Spearman correlation coefficient and p-value for all non-chr21 genes.

(B) Individual significantly-enriched terms (combined p ≤ 0.1) from each of the categories in Fig. 4(D,E), selected by a concordance measure (euclidean distance from origin in D). Colors denote NES, bubble size the individual -log10(p.adj).

**Figure S4**

Comparison of terms over-represented in GSEA across hiPSC T21, neural T21, T21 dosage correction (“rescue”), and the dox-effect in non-transgenic hiPSCs. Terms from Hallmark, KEGG, and WikiPathways (WP) are shown. All terms are sorted by hiPSC T21 NES (normalized enrichment score from GSEA), with a p.adj thresholds of p ≤ 0.1 for hiPSC T21, and additionally p.adj ≤ 0.2 for the three additional panels (neural T21, T21 rescue, and dox-effect).

**Figure S5**

(A) Representative images of TS21 interneuron and euploid control cultures transduced with hSyn1:RGEDI-P2a-EGFP lentivirus. Scale bar = 100 μm.

(B) Quantification of the GEDI ratio (mean RGEDI/mean EGFP) across each interneuron per time point (T21 n= 8810, D21/euploid control n= 15,999). Live/dead threshold of 0.15 (red line)

(C) as in (B), but across each forebrain neuron per time point (T21 n= 30,735, D21/euploid control n= 24,950).

(D) Number of cells tracked by segmentation in independent wells of T21 and D21 neurons across all timepoints.

(E) Fluorescence intensity signal of RGEDI, EGFP and RGEDI/EGFP ratios in live (RGEDI/EGFP ratio < 0.15) T21 and euploid control neurons (p ≤ 1×10^8^, linear mixed model with Holm correction).

(F) Anti-GFAP (red) and DAPI (blue) staining of cultures from (A) showing abnormal glia morphology. Scale bar = 300 μm.

**Figure S6**

(A) S100B+ cell density and representative images of E18.5 wildtype and Ts65dn cortical slices (100 μm scale bars). Two-tailed, independent samples Student’s t-test p ≤ 0.05 (*).

(B) Ki67^+^ cell density and representative images of E18.5 wildtype and Ts65dn cortical slices (60 μm scale bars). Two-tailed, independent samples Student’s t-test p > 0.05 (n.s.).

**Figure S7**

(A) Sample proportions (% nuclei) of assigned cell types, cell cycle stage and *XIST* status faceted by cell type group.

(B) Paternal allele fraction by cell, aggregated over all chr21 genes, and faceted by cell type group. Cells split by sample (D21a/b, T21XIST-nodox, d0+dox, pD+dox) and *XIST* status (0, >1 counts denoted “X-“ and “X+”), except for euploid lines (absent XIST transgene, grey). Median paternal allele fraction and significance of difference in medians relative to T21-nodox cells (Wilcoxon p-value) are denoted below and above boxplot, respectively.

(C) Left: Normalized gene expression ratio (chr21 over genome mean excluding chr21) by cell, aggregated over all chr21 genes, and faceted by cell type group. Cells are split as in B, with median expression ratio and difference in medians relative to T21-nodox cells (Wilcoxon p-value) denoted above and below boxplot, respectively. Right: empirical cumulative distributions (eCDF) of corresponding panels on the left, colored by sample and XIST-positive cells represented by dashed lines. T21-nodox 95% confidence interval denoted by transparent ribbon (purple). Sample-colored p-values (Kolmogorov-Smirnov, KS-test) denotes significance of distribution of shift from the T21-nodox sample. Grey lines indicate the 50^th^ percentile cell (horizonal), and chr21 to genome-wide mean ratio (vertical 1.0 and 1.5).

(D) Left: Normalized expression ratio by chr21 gene, aggregated over cells, and faceted by cell type group. Mean ratios per gene calculated from cells are split as in C. Respective median expression ratios and difference in medians relative to T21-nodox cells (Wilcoxon p-value) denoted above and below boxplots. Right: eCDF of corresponding panels of the left, with lines colored as in C. T21-nodox confidence-interval, sample-colored p-values (KS-test) and grey vertical lines s in C, with the horizontal grey line indicating the 50^th^ percentile chr21 gene.

(E,F) Cumulative distributions aggregating across cell type groups (right panels in C & D), excluding the “Other” group. Lines, sample-colored p-values (KS-test), grey horizontal and vertical lines as in C and D, respectively.

(G) Differential expression of chr21 genes relative to XIST-negative postDAPT and T21-nodox cells. Average log2(FC) values in euploid cells (x-axis) correlate with XIST-positive d0 and pDAPT log2(FC) values (y-axis), faceted by cell type group. Colors and transparency denote respectively, ROC power and ROC power of the euploid relative to T21 comparison. Regression line (black), pearson coefficient and p-value (Fisher-transformed) denoted on each plot.

**Figure S8**

(A) SC-transformed sample-averaged log2(FC) values relative to XIST-negative T21-nodox and pDAPT cells faceted by cell type group. Median sample-averaged log2(FC) values of euploid (D21a/b), and XIST-positive d0+dox and pD+dox samples, and significance of median differences (Wilcoxon p-value) from T21 to euploid comparison denoted below and above corresponding boxplot.

(B) Loess-smoothened SC-transformed log2FC values of differential expression comparisons (as in A, with 95% confidence intervals) across chr21 coordinates of transcriptional start sites (TSS, in Mbp).

(C) SC-transformed sample-averaged log2(FC) values of d0+dox and pD+dox comparisons (colored as in A,B) plotted over the TSS distance from the *XIST* transgene integration site, faceted by cell type group. Regression lines, 95% confidence interval ribbons, pearson coefficient (R), and p-value (fisher-transformed R) indicated on each plot, black horizontal line denotes no change.

(D) Normalized enrichment scores (NES, x-axis) of GSEA against Hallmark and Canonical pathways MSigDB subsets in euploid (center) and XIST-positive d0+dox and pD+dox cells. Plotted categories (y-axis) limited to significant enrichment across at two comparisons (pairwise Fisher’s method combined p-value ≤ 0.005). Colors and transparency denote respectively, NES sign and – log10-transformed q-values (capped at 3, q <= 0.001).

## MATERIALS & METHODS

### EXPERIMENTAL DESIGN

#### Reprogramming, hiPSC validation and culture

Human male fibroblast line (GM00260) mosaic for trisomy 21 (T21) was obtained from the Coriell Institute (Camden, NJ) and reprogrammed by the University of Connecticut’s Cell and Genome Engineering Core, using the CytoTune hiPSC 2.0 Sendai Reprogramming kit (Thermo Fisher Scientific, Waltham, MA). Genomic DNA from hiPSC clones was screened via duplex Taqman Copy Number (CN) Assay (*DYRK1A* – Hs01748441_cn and RNaseP, Thermo Fisher Scientific) to select euploid and T21 lines, followed by additional screening for Chr17q Taqman CN assay (*SEC14L1*, Hs05476397_cn), and the hPSC Genetic Analysis Kit (Stemcell Technologies, Vancouver, Canada). Interphase and M phase nuclei were also screened via FASTFISH (WiCell, Madison, WI) using Chr 20q (BCL2L1) and centromere enumeration probes for Chr 8, 12, 17, and X, alongside a SureFISH assay for Chr 21q (DSCR8). Selected hiPSC lines were further screened by aCGH (Infinium BeadChip CytoSNP 850K assay, Illumina, San Diego, CA) at the Center for Genome Innovation, Institute for Systems Genomics at the University of Connecticut.

T21 and euploid hiPSCs were cultured on mitotically inactive mouse embryonic fibroblasts (mEFs) in standard hiPSC media in 5% CO2 at 37°C (80% DMEM/F12, 20% Knockout Serum Replacement, 1% Glutamax, 1% Non-Essential Amino Acids, 0.1% ß-Mercaptoethanol, and 8 ng/mL bFGF, Thermo Fisher Scientific), and passaged weekly by manual disruption of 6-10 representative colonies. For hiPSC cultures requiring doxycycline induction, 2 μM doxycycline was added to the culture media.

#### Plasmid construction

For PCR amplifications, either Herculase II polymerase (Agilent, Santa Clara, CA), Platinum Taq High Fidelity (Thermo Fisher Scientific), Advantage 2 polymerase (TaKaRa Bio, Shiga, Japan), or Q5 Hi-Fidelity DNA Polymerase (New England Biolabs) was used. For cloning reactions except the large *XIST* insert steps, ligations were performed with T4 DNA ligase, 2,000,000 units/mL (New England Biolabs) and DH10B homemade chemically competent cells were used at 37°C. For all cloning with XIST cDNA fragments aside from ligation of oligo not followed by transformation (which used the same T4 DNA ligase), the TaKaRa DNA Ligation Kit LONG (TaKaRa Bio) was used along with NEB Stable Competent E. coli (New England Biolabs) with bacterial incubations at 25-30°C to mitigate instability in initial steps observed at 37°C.

To clone the XIST transgene cDNA in a temporary pUC19 vector, a 10 kb fragment containing most of XIST exon 1 was cut from a fosmid (WI2-935P22, BACPAC Genomics) with SacII and PshAI. Linker oligos were used to clone the 10 kb SacII and PshAI fragment into the KpnI and XbaI sites of pUC19, respectively. The 5’-most end of *XIST* exon 1 was PCR amplified, concurrently adding a 5’ PmeI restriction site. The resulting PCR product was digested with PmeI and BtgZI and cloned in between the PmeI/BtgZI restriction sites of the XIST exon 1-containg plasmid. The 3’ end of exon 1 through exon 6 was amplified from cDNA of Wi-38 cells (AG07217, Coriell Institute), while adding a 3’ SalI restriction site. The resulting PCR was digested with PshAI and SalI and cloned between the PshAI and SalI sites of the plasmid from the prior step to complete the XIST cDNA construct. The *UBC* promoter from the FUGW construct ^79^ (Addgene plasmid #14883) was cloned into AAVS1-Neo-M2rtTA ^80^ (Addgene plasmid #60843). The *UBC* promoter was PCR-amplified with flanking PvuII and FseI sites, and cloned into the AleI and FseI sites on the AAVS1-neo-M2rtTA plasmid. This rtTA-advanced ORF was replaced with rtTA3G from the pCMV-Tet-3G plasmid (TaKaRa Bio), by XbaI and XmaI digestion and cloning of three separate fragments to create the final rtTA vector AAVS1-NEO-UBrtTA-3G.

To generate the final targeting vector, pTRE3G (TaKaRa Bio) was used as the backbone, following excision of an XbaI fragment to reduce the size of the vector via removal of the SV40 small intron, NLS, and SV40 polyA sequences. The 5’ homology arm for chr21 was generated by PCR with EcoRI sites on each primer using the genomic DNA of the destination cell line as the template. This PCR product was ligated into the EcoRI sites of the pTRE3G vector to make pTRE3G-5P. Next a fusion PCR product was generated, harboring a BGH polyA amplified from pl452 ^81^ and the 3’ homology arm using overlapping PCR from the genomic DNA of the destination cell line. This PCR product was cloned into the PstI and BamHI sites of the pTRE3G-5P to make pTRE3G-5P3P. Next a loxP-flanked PGK-puro-TK fragment was removed from vector pl452-puroTK by XhoI digest. The fragment was cloned into the XhoI sites of the vector pTRE3G-5P3P. This pl452-puroTK vector was derived from plasmids pl452 and plasmid pLCA.66/2272 ^82^ (Addgene plasmid #22733) by removing the PGK-puroTK cassette with AgeI and BbsI and cloning that fragment into pl452 using the same enzymes. To add the *XIST* cDNA construct, this CMV-puroTK-TRE3G vector was digested with MluI and SbfI, and a phosphorylated linker oligo was ligated to purified, digested backbone, creating a SacI-compatible overhang. Following purification, this vector was ligated to the SbfI-SacI fragment containing the *XIST* cDNA construct, to create the TRE3G-XIST cDNA targeting vector.

#### hiPSC targeting and screening

Targeting was performed using a Lonza Nucleofector 4D (Lonza, Basel, Switzerland) with primary cell kit P3 and program CD-150. AAVS1-Neo-ubrtTA was targeted to chr21 of the T21 cell line 198-5. Two μg of each AAVS1-targeting TALEN ^83^ was nucleofected along with 6 μg of AAVS1 integration vector. Cells were plated on mitotically inactivated DR4 MEFs (ATCC, Manassas, Virginia) post-nucleofection. After 96 hours, cells were cultured with G418 at a concentration of 25 μg/ml. Selection was done for a total of 6 days. Resulting colonies were isolated and screened for targeting. cDNAs from targeted clones were tested for expression of rtTA by RT-qPCR. AAVS1 TALEN-L (Addgene plasmid #59025) and AAVS1 TALEN-R (Addgene plasmid #59026) ^84^ were used for targeting the UBrtTA . Integrants were screened using (AAVS1 scr F/Neo R and AAVS1 3’scr R/rtTA F). For targeting XIST transgene via CRISPR, guides designed to target the chr21P copy without cutting the chr21^M/M2^ alleles were cloned into PX459V2 (Addgene plasmid #48139) ^85^ at the BbsI site by annealing and ligating overlapping oligos. Two μg each this CRISPR guide plasmid was used along with 6 μg XIST cDNA targeting vector. Cells were plated on DR4 MEFs and after 96 hours, puromycin was added to the culture at a concentration of 0.5 μg/ml for a total of 4 days. Resulting colonies were isolated and genotyped for targeting by PCR. Clones were verified by sequencing across the target to verify the loss of the targeted allele.

#### Nucleic acid purification

Genomic DNA was purified from fresh or frozen pelleted cells using three different methods depending on application. For colony PCR screening, genomic DNA was extracted via standard high-salt chloroform extraction. For standard PCR, and array hybridization, cell pellets were resuspended in either cell lysis buffer 1 (10 mM Tris-HCl, pH 8.0; 100 mM NaCl; 10 mM EDTA, pH 8.0; 1% SDS; 200 μg/ml Proteinase K), or cell lysis buffer 2 (10 mM Tris-HCl, pH 8.0; 200 mM NaCl; 10 mM EDTA, pH 8.0, 0.5% Sarcosyl; 1 mg/mL Proteinase K) and incubated at 55°C for 3 hours to overnight with optional RNAse A, 10 μg/mL treatment following at 37C for 5 minutes. Genomic DNA was isolated using Phenol:Chloroform:Isoamyl Alcohol (25:24:1, v/v) (Invitrogen, Waltham, MA), precipitated with either isopropanol or 300 mM sodium acetate with 2 volumes of 100% chilled ethanol and 15 minutes at -80C, washed with 70% ethanol, and resuspended in TE buffer. For linked-read WGS, high molecular weight (HMW) genomic DNA was prepared from fresh or frozen hiPSC cell pellets, following the protocol listed in the Chromium Genome Reagent Kits v2 User Guide (10x Genomics, Pleasanton, CA), using the MagAttract HMW Kit (Qiagen, Germantown, MD) with minor modifications. Plasmid and fosmid DNA was purified with either the Zyppy Plasmid Miniprep kit or ZymoPURE II Plasmid Midiprep kit (Zymo Research, Irvine, CA). Following restriction digests or PCR, DNA purification was performed with the Monarch PCR & DNA Cleanup kit (New England Biolabs, Ipswich, MA), or the DNA Clean & Concentrator Kit (Zymo Research). For RNA isolation, the PureLink RNA Mini Kit (Invitrogen, Carlsbad, CA) was used, followed by cDNA synthesis using the iScript gDNA Clear cDNA Synthesis Kit (Bio-Rad Laboratories, Hercules, CA).

#### Quantitative RT-PCR

Gene expression was measured via quantitative real-time PCR (qRT-PCR) using the iTaq Universal SYBR Green Supermix (Bio-Rad Laboratories) and the ΔCt method. Primer specificity was confirmed by melting curve analysis, and relative expression was calculated as %GAPDH (100/2^ΔCt^). Taqman CN assays were normalized to 198-1 genomic DNA. HOT FIREPol Probe Universal PCR Mix Plus – no ROX (Solis BioDyne, Tartu, Estonia) was used per manufacturer’s instructions, using 4 ng of high-quality genomic DNA template in 20 μl reactions. The hPSC Genetic Analysis Kit (Stemcell Technologies) was used following the manufacturer’s instructions. All qPCR reactions were performed in triplicate in a Bio-Rad CFX 96 Real-Time System.

#### Cytogenomics, DNA methylation, Linked-read WGS and mRNA sequencing

For cytogenomic analysis, genomic DNA was labeled and hybridized to the Infinium BeadChip CytoSNP 850K array (Illumina, San Diego, CA) according to the manufacturer’s protocols. For DNAme analysis, genomic DNA was bisulfite-converted using the EZ DNA Methylation Kit (Zymo Research, Irvine, CA), then labeled and hybridized using the Infinium Methylation EPIC BeadChip Kit (Illumina) following standard protocol of each manufacturer. Both types of arrays were scanned on a NextSeq 550 system (Illumina). HMW genomic DNA was sequenced to ∼30x coverage on a NovaSeq 6000 (Illumina), following library preparation on the 10X Genomics Linked-Read platform at the Center for Genome Innovation, Institute for Systems Genomics at the University of Connecticut. For standard mRNA-seq, libraries were prepared using the Illumina stranded mRNA Kit and 100bp paired-ends reads were sequenced to an average depth pf 40 million reads/replicate on the NovaSeq 6000 (Illumina, San Diego, CA).

#### Immunofluorescence and XIST RNA-FISH

For *XIST* RNA FISH, cells were permeabilized with 0.5% Triton-X and 2mM ribonucleoside vanadyl complex (RVC) in CSK Buffer (10mM PIPES pH 6.8, 100mM NaCl, 3mM MgCl2, 300nM sucrose), fixed in 4% PFA in PBS, then dehydrated through a series of ethanol washes (70%,80%, 90%, and 100%). Digoxigenin-labeled *XIST* probe hybridization was performed overnight in a humidity chamber at 42°C. Anti-digoxigenin antibody (ThermoFisher; 700772) was hybridized for 1 hour at RT. Secondary antibody hybridization (Alexa Fluor; Invitrogen; 1:500) followed for 1 hour at RT. Nuclear DNA was counter-stained with Hoechst 33342 dye. Slides were imaged with an EVOS FL Auto2 system (Thermo Fisher Scientific).

For immunofluorescence cells were grown in Nunc Lab-Tek II glass chamberslides coated with poly-d-lysine and laminin (for neural differentiation) or permanox chamberslides with MEFs (iPSCs). Cells were fixed in 4% paraformaldehyde (Electron Microscopy Sciences, Hatfield, PA) for 10 minutes at room temperature (RT), rinsed with PBS 3 times for 5 minutes and then permeabilized in 1% Triton-X in PBS for another 10 minutes. Samples were blocked in blocking buffer containing PBS, 0.1% Triton-X, 5% BSA and 2% normal goat serum (MilliporeSigma). Cells were placed in antibodies diluted in blocking buffer overnight at 4°C in a humidity chamber. Primary antibodies were used as follows: MAP2 (Abcam; ab5392; 1:10,000), S100B (Sigma; s2532; 1:1000), H2AK119ub (Cell Signaling; 8240, 1:1600). Following washes in 0.1% Triton-X in PBS, 3 times for 5 minutes, cells were labeled with Alexa Fluor secondary antibodies at 1:500 for 1 hour at RT. Nuclei were counterstained with Hoechst 33342and slides preserved in Prolong Gold Antifade Mountant (Thermo Fisher Scientific).

#### Neural differentiation

T21 and euploid hiPSCs were passaged manually onto mEFs and in hiPSC medium. On day 2 post-passage, cultures were switched into N2B27 neural differentiation medium (Neurobasal medium (Gibco), 2% B27 (Gibco), 1% N2(DMEM:F12 (Gibco), 25 μg/ml insulin (Sigma), 100 μg/ml holo-transferrin, 100 μM putrescine, 20nM progesterone, 30nM selenite), 1% ITS-A (Gibco), 1% Penicillin-Streptomycin (Gibco), and 1% Glutamax (Gibco), supplemented with noggin at 500 ng/ml (R&D Systems). Cultures were fed and supplemented with Noggin every other day through day 10. For neural differentiation with doxycycline induction, 1 μM doxycycline, or 0.01 μM for low-dose cultures, was added to the culture media. Cultures were fed N2B27 until the appearance of mature rosette structures (D14-D16). Rosette-stage cultures were passaged manually 1:1 onto poly-d-lysine (PDL) and laminin-coated tissue culture plates and grown to confluence. Cells were cultured until d23 and passaged enzymatically with Accutase (Invitrogen) onto PDL and laminin-coated substrates at 1.9×10∧4 cells/cm^2^-2.2×10∧4 cells/cm^2^. At d25, cultures were switched to Neural Differentiation medium (NDM) (Neurobasal medium (Gibco), 2% B27 (Gibco), 1% non-essential amino acids (NEAA; Gibco), 1% Penicillin-Streptomycin (Gibco) 2mM Glutamax (Gibco) 10 ng/ml brain-derived neurotrophic factor (BDNF; Peprotech), 10ng/ml glial-derived neurotrophic factor (GDNF, Peprotech), 1 μM cyclic AMP (cAMP, Sigma), and 200 μM ascorbic acid (AA, Sigma). Cells were cultured in NDM until assay endpoint. For DAPT-treated cultures, DAPT (Tocris Bioscience) was added for 5 days at 10 μM. For EdU labeling, Edu was added at 5 μM for 24 hours, and detected using the Click-iT EdU Imaging Kit (Invitrogen). Doxycycline (1 μM) was added to EdU-labeled, pDAPT cultures at the time of EdU addition/labeling.

For longitudinal imaging of forebrain cortical neurons and interneurons, euploid (198-1) and T21 (198-5) hiPSCs were maintained on Matrigel coated plates in mTESR media before differentiation. Forebrain cortical neurons were derived from hiPSCs as described in ^86^. Briefly, neural differentiation was induced with Smad inhibitors Dorsomorphin (1.5 μM) and SB431542 (10 μM) for 11 days. Inhibitors were removed on day 12, and supplemented with retinoic acid (0.05 μM) on day 16. At day 20, cells were dissociated with 0.05% trypsin-EDTA, resuspended in media with BDNF (2ng/mL), GDNF (2ng/mL) and retinoic acid (0.05 μM) and plated 1:2 onto Poly-L-ornithine-Laminin coated dish. For forebrain interneuron differentiation, hiPSCs were first dissociated as colonies with CollagenaseIV/Dispase (1mg/mL each; Invitrogen) and further into single cells using 0.05% trypsin-EDTA. Then, 10,000 cells were plated on low attachment round bottom 96-well plate to form suspension embryoid bodies (sEB) in optimized B27 + 5 factor (B27+5F) media (as described in ^87^). On day 7, sEBs (35–40 per well) were transferred to 6-well Matrigel coated plates, without ROCK inhibitor. On day 14, media was replaced with the addition of Purmorphamine (1μM) only. On day 25, adherent embryoid bodies (aEB) were dissociated using 0.05% trypsin-EDTA, passed through 70 μm cell-strainer, and plated at a density of 10,000-25,000 cells/cm^2^ onto Poly-L-ornithine-Laminin coated plates. Neurons were matured in B27 media containing DAPT (10 μM) and BDNF (2ng/mL). The pHR-hSyn1-RGEDI-P2a-EGFP lentivirus (MOI=1) was added to forebrain cortical neurons (Day 35) and interneurons (Day 56) for survival analysis as described ^57^ and imaged with robotic microscopy ^88^ every 12 hours for 10 days. The cells were maintained in an environmental chamber with 5% CO2 during imaging. These live cell images were taken on an inverted microscope (Nikon Ti-E) at 20X magnification. A customized software controlling Micromanager (version 2.0 gamma) is used to allow the automated runs return to the same field of view on the plate for longitudinal imaging in 3D space.

#### Single-nuclei RNA sequencing

Neural cultures were treated with TrypLE (Life Technologies) for 4 minutes at 37C, and cells counted with Trypan blue on the Countess Automated Cell Counter. Nuclei isolation was performed with the Nuclei Isolation Kit: Nuclei EZ Prep (Sigma-Aldrich) with some modifications. Briefly, cells were centrifuged at 1330 rpm for 3.5 minutes, media aspirated, and resuspended in Nuclei PURE Lysis Buffer containing 1M DTT, 10% Triton X-100 and RNAsin. Nuclei were centrifuged at 500g and 4C for 5 minutes in a swing-bucket rotor centrifuge, and lysis buffer was aspirated. Nuclei resuspended in Nuclei PURE Sucrose Cushion buffer and RNAsin were pelleted at 500g and 4C for 5 minutes, resuspended again in 25% iodixanol solution (0.28 ml of Nuclei PURE Sucrose Cushion buffer containing RNAsin + 0.2 ml of 60% iodixanol), and under-layered by corresponding 30% and 35% iodixanol solutions. Nuclei traverse the 25% and 30% layers of this density gradient upon centrifugation (3166g at 4C for 30 minutes), and were collected from the 30% to 35% interface in volume of 0.3 ml. Resuspended in PURE Storage Buffer (1 ml) and (5 μl) of RNAsin, nuclei were pelleted at 3166g for 15 minutes at 4C, resuspended in remaining 0.1 ml, and counted again. Isolated nuclei were stored at -80 until fixation. Nuclei fixation was performed according to the Evercode Fixation v2 protocol (Parse Biosciences). Nuclei were counted after fixation and barcoded using the Evercode WT mini v2, following the manufacturer’s instructions. Sublibraries were quantified using the Agilent Tapestation, pooled in proportion to their respective nuclei counts, and sequenced on MiSeq v2 and NextSeq 550 v2.5 mid-output kits (Illumina).

#### Animals

Ts(1716)65Dn (Ts65Dn; strain #005252) mice were obtained through crossings of a B6EiC3Sn a/A-Ts (1716)65Dn (Ts65Dn) female to F1 hybrid B6C3F1/J males purchased from The Jackson Laboratory. Genotyping was performed by amplifying genomic DNA with an in-house protocol. The colony of Ts65Dn mice was maintained in the Animal Facilities in the Barcelona Biomedical Research Park (PRBB, Barcelona, Spain). Mice received chow and water ad libitum in controlled laboratory conditions with temperature maintained at 22 ± 1oC and humidity at 55 ± 10% on a 12h light/dark cycle (lights off 20:00h). The control group consisted of wild-type (WT) euploid littermates. All experimental procedures were approved by the local ethical committee (Comité Ético de Experimentación Animal del PRBB (CEEA-PRBB); procedure number MDS-12-1464P3), and met the guidelines of the local (law 32/2007) and European regulations (EU directive n° 86/609, EU decree 2001-486) and the Standards for the use of Laboratory Animals n° A5388-01 (NIH).

#### Slice immunohistochemistry

Breeding pairs were established so that vaginal plugs could be checked twice daily. The presence of a vaginal plug was designated as E0.5. A 10% weight gain at E10 was used to confirm pregnancy. Male and female embryos were collected from pregnant trisomic dams at E18.5, chilled, decapitated immediately and processed for fluorescent immunohistochemical staining. Fetal heads were immersed in Paraformaldehyde 4% (PFA) (Sigma Aldrich) and the brain was dissected 24 hours later and immersed again in PFA o/n. Brains were cryoprotected by immersion in 30% sucrose solution and embedded in OCT for posterior histological processing. Embryos heads were embedded in optimal cutting temperature compound (OCT) and embedded tissue was frozen rapidly and either stored at − 80 °C or immediately sectioned into 20-μm thick frozen sections using a cryostat (Leica CM3050S). Histological sections were placed in Superfrost® slides and placed at -20°C. We processed two sections at three different coordinates (+1.81/2.52/3.15 from Bregma). Slices were fixed with methanol and incubated with blocking buffer (Tris-Buffered Saline, TBS, with 0,5% Triton-X 100 and 6% normal donkey serum). Primary antibodies were diluted in TBS ++ (TBS with 0,1% Triton-X 100 and 6% normal donkey serum): anti-S100b (Rabbit, 1:1000; Synaptic systems, #287003) and anti-Ki67 (Rabbit, 1:500; Abcam, ab15580). Slices were incubated with primary antibodies overnight. After three washes with TBS, sections were incubated with the following secondary antibodies diluted in TBS ++: goat anti-rabbit (1:500; Alexa 594, Invitrogen A11012) and goat anti-rabbit (1:500; Alexa 488, A-11008) . Slices were washed with TBS three times and incubated ten minutes with Hoechst 33342. Finally, sections were mounted on glass slides using an aqueous mounting media, Mowiol. To quantify the density of S100B cells six sections per animal were imaged using a Leica TCS SP5 inverted scanning laser microscope (Leica Microsystems). 15 – 16-μm thick z-stacks (512 x 512 resolution, 8 bits) of each region of interest (ROI) in the cortex of embryos, were acquired using a 20X objective (N.A = 0.8).

### STATISTICAL ANALYSIS

#### Image analysis

For IF (H3K27me3 and H2AK119ub) and *XIST* FISH in hiPSCs, automated image quantification relative to all Hoechst33342 labeled cells was performed using a custom CellProfiler pipeline ^89^, which was run in 2 phases. First, images were optimized for computer recognition using the following CellProfiler functions on each channel of each image: RescaleIntensity, CorrectIlluminationCalculate, CorrectIlluminationApply, and ReduceNoise. Then signals for nuclei, IF, and *XIST* FISH were quantified using the IdentifyPrimaryObjects functions, and counts for IF or *XIST*-positive nuclei were obtained using RelateObjects and FilterObjects. Images were output alongside outline annotations for objects recognized by CellProfiler to visually verify accuracy and consistency of automated counts. For EdU and *XIST* FISH, as well as IF of hiPSC-derived neural lineages (MAP2, S100B), images were imported and labeled cells counted manually using ImageJ (NIH, Bethesda). Differences in median fraction of labeled cells were assessed by two-tailed Wilcoxon rank-sum test.

For longitudinal imaging of forebrain cortical neurons and interneurons, images were segmented using EGFP morphology, background subtracted, aligned together across timepoints, and then extracted channel intensity and feature information from each neuron into a csv file. Survival analysis was performed by defining the time of death as the point at which the GEDI ratio (RGEDI signal: EGFP expression) of a longitudinally imaged neuron exceeds the empirically calculated GEDI threshold ^57^. Additionally, single cells were tracked ^58^ and cumulative death was calculated using Linear mixed model with Holm correction in R studio (version 4. 0. 4).

For slice immunochemistry images, labeled cells were manually counted using ImageJ (NIH, Bethesda). Cell counting was performed using the Cell Counter plugin on ImageJ in a maximum projection of all z-stacks (1 μm step size). Cell counts from the cortex were processed first by image, then by coordinate (two images were taken from each coordinate), then by animal and finally by genotype. To quantify the density of Ki67 cells three sections per animal were imaged in the same form than S100B but imaging the ventricular zone of the cortex, the area with higher proliferation. An area of 100 µm^2^ was delimited to normalize across sections and due to the high density of cells in this area. Outliers in data sets from both euploid and trisomic mice were statistically determined using a calculation of interquartile range (IQR). All data points outside of the IQR fences were excluded from analysis without bias. All variables were assessed with a two-tailed, independent samples Student’s t test, and passed both the Shapiro-Wilks normality test and an equal variance test.

#### Expression and DNA methylation analysis

FASTQ files were quality controlled using FASTQC, reads were filtered and adapter-trimmed was done using BBDuk ^90^. Expression quantification was performed using Salmon ^91^, and differential expression analysis was performed using DESeq2 ^92^. Gene set enrichments were performed using the GSEA function from the R package clusterProfiler ^93^ on the following gene sets: Biocarta, GO (Biological Processes, Cellular Components, and Molecular Functions), KEGG, Pathway Interaction Database, Reactome, and WikiPathways. For DNAme analysis, the MethylEPIC IDAT files were processed using the R package minfi and its preprocessIllumina function to obtain methylation β-values ^94^. Promoter probes were defined as having one of the following designations in the array manifest file: TSS1500, TSS200, or 5_UTR.

#### Phased allelic RNA-seq analysis

The 10X Genomics Long Ranger pipeline was used to align reads, and call and phase SNPs, indels, and structural variants from linked-read WGS reads. The processed FASTQs from above were aligned using HISAT2 to the hg38 human reference genome ^95^. Relevant chr21 SNPs were extracted from the Long Ranger output and used to get haplotype gene expression counts from aligned RNA-seq bam files using the phASER software package ^96^. While this provided reliable intragenic phasing, for intergenic phasing on chr21 we relied on the imbalanced haplotype gene expression in the T21, compared to its isogenic euploid controls. The higher expressed haplotype was designated as the major (duplicated or chr21^M/M2^) haplotype and the other as the minor haplotype (chr21^P^).

#### Single-nuclei RNA-seq analysis

FASTQ files were processed using the split-pipe pipeline (Parse Biosciences), and imported to Seurat ^97^. We excluded nuclei with over 10% reads mapping to the mitochondrial genome, fewer than 2500 mapped reads, fewer than 1500 genes detected, or over 7500 genes detected. Following normalization and cell cycle scoring, the data were SC-transformed ^62^ while regressing out the mitochondrial read fraction, and samples integrated. UMAPs were projected using the first 30 principal components, and clusters labeled under default settings. Cell types were identified and grouped using markers from ^60^ and sctype ^61^. Allelic counts from single nuclei were phased as described above, and aggregated across all chr21 genes to mitigate the impact of transcriptional bursts. Chr21 dosage by cell (aggregating across chr21 genes), or by gene (aggregating across cells) was performed on counts-per-million (cpm) normalized expression. Differences in median LAF or chr21 dosage ratio were assessed by two-tailed Wilcoxon rank-sum test, and differences in distribution by two-sample Kolmogorov-Smirnov test.

For differential expression analysis, Seurat functions PrepSCTFindMarkers and FindMarkers using the ROC (receiver operating characteristic curve) test were applied to compare euploid, *XIST^+^* d0 and pD cells to *XIST-*negative T21 cells by cell type group. GSEA was performed as for bulk RNA-seq while limiting gene terms to Hallmark and canonical pathways (KEGG, Reactome, wikipathways, PID, biocarta) of MSigDB.

## ACKNOWLEDGEMENTS

This work was supported by grants from the LuMind IDSC Foundation and the NIH (R35GM124926, with additional support from R01HL141324) to S.F.P . We are grateful to Gordon Carmichael and James Li for helpful comments on our manuscript. We also thank the University of Connecticut’s Cell and Genome Engineering Core and Center for Genome Innovation for reprogramming, as well as mRNA-seq, CytoSNP, linked-read WGS and methylEPIC core services, respectively. Support for work conducted at the Gladstone institutes (S.F.) came from R01AG064579 with additional support from R01LM013617, RF1NS128800 and the JSRM Foundation. Work at the Centre for Genomic Regulation (M.D.) was supported by the Fondation Jérôme Lejeune #2002, Fundació La Marató De TV3 202212-30-31-32, and Agencia Estatal de Investigación (PID2019-110755RB-I00/AEI / 10.13039/501100011033). We acknowledge support of the Spanish Ministry of Science and Innovation to the EMBL partnership, the Centro de Excelencia Severo Ochoa and the CERCA Programme / Generalitat de Catalunya, and the FPU fellowship (FPU19/04789) from Ministerio de Universidades (to M.S.N.) The CIBER of Rare Diseases is an initiative of the ISCIII.

## DECLARATION OF INTERESTS

No competing interests

## Notes

### Competing Interest Statement

The authors have declared no competing interest.

### Summary of Updates

Additions to references, materials and methods, minor changes to the main text, acknowledgments, and figure legends.

