## Supplemental Figures for "A dynamic *in vitro* model of Down Syndrome neurogenesis with Trisomy 21 gene dosage correction"

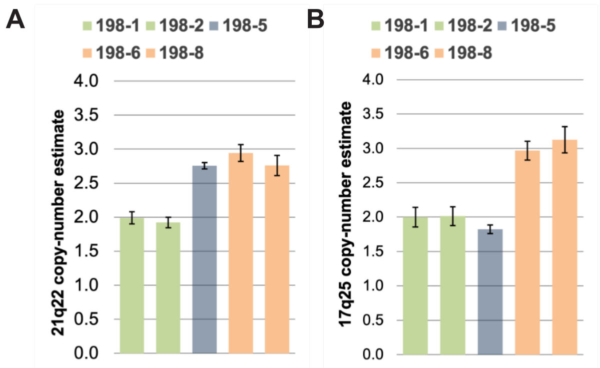


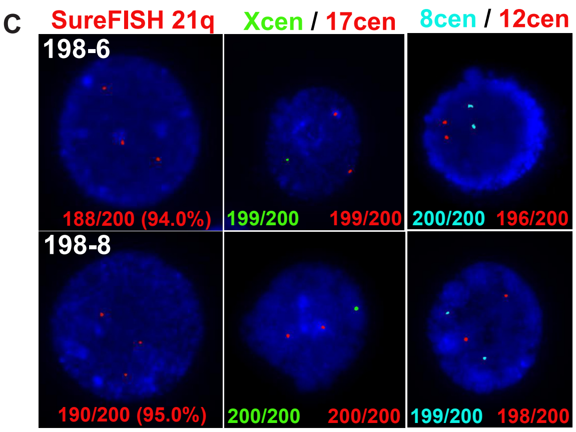


D

| CNV type: chr. band (size) | GM 00260 | 198-1 | 198-2 | 198-6 | 198-8 | 198-5 | rTTA 5-1 | rTTA 5-9 | rTTA 5-10 |
|---|---|---|---|---|---|---|---|---|---|
| Loss: 1p13.2 (439 kb) | | | | | x | | | | |
| Loss: 2q23.3 (439 kb) | | | | x | | | | | |
| LOH: 3p21.2-3p21.31 (3.9 Mb) | x | x | x | x | x | x | x | x | x |
| Gain: 4q25 (738 kb) | | | | x | | | | | |
| Gain: 7q11.21 (650 kb) | x | x | x | x | x | x | x | x | x |
| Gain: 8q13.2-q13.3 (497 kb) | | | | | x | | | | |
| Loss: 11q14.2 (791 kb) | | | | | x | | | | |
| Gain: 17q21.2-17q25.3 (42 Mb) | | | | x | x | | | | |
| Loss: 18q22.1-18q22.2 (579 kb) | | | | x | | | | | |
| Trisomy: 21p11.2-21q22.3 | x | | | x | x | x | x | x | x |

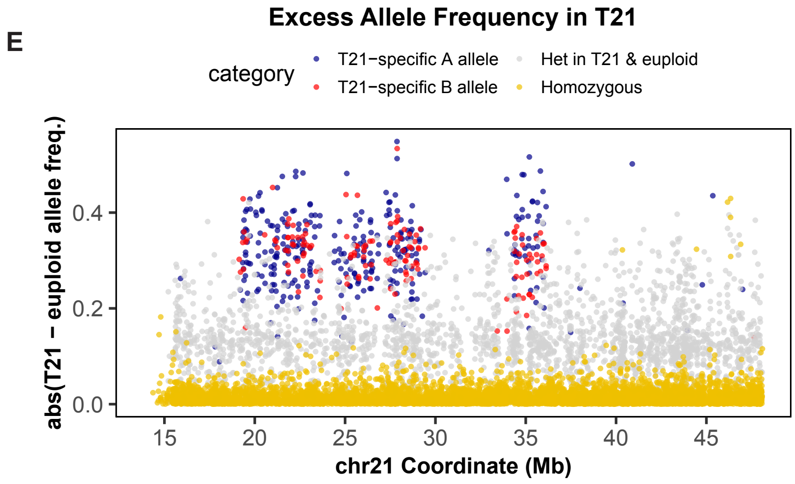


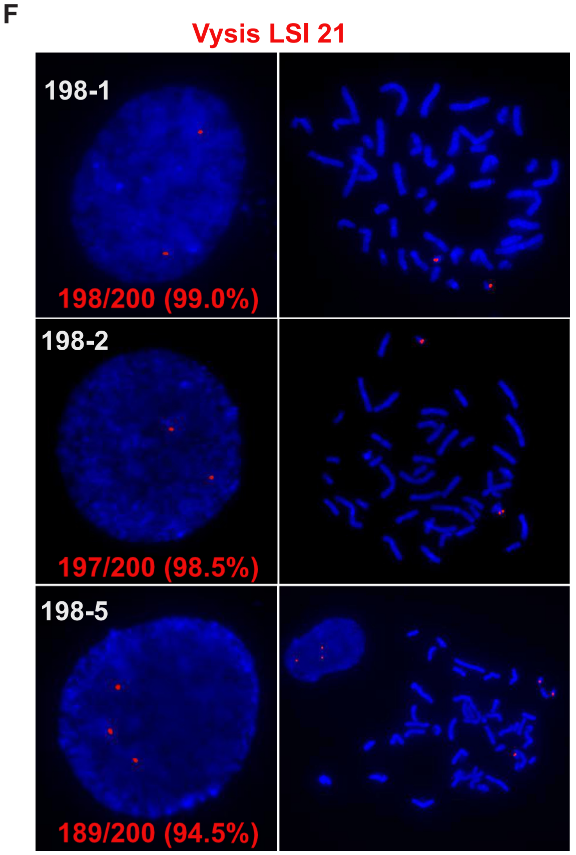


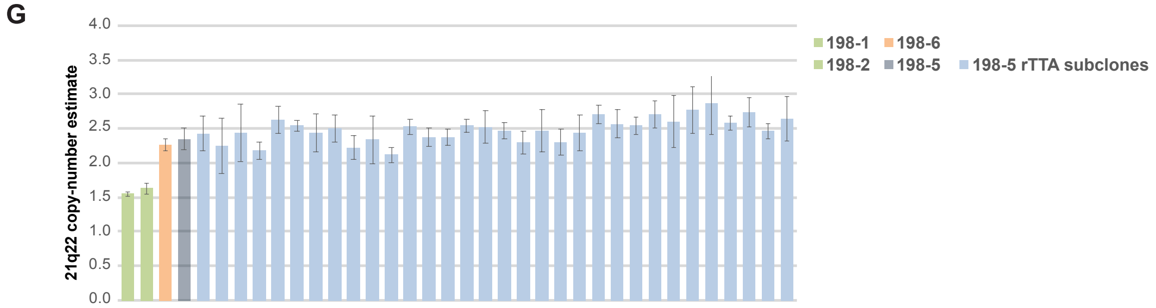


**Figure S1**

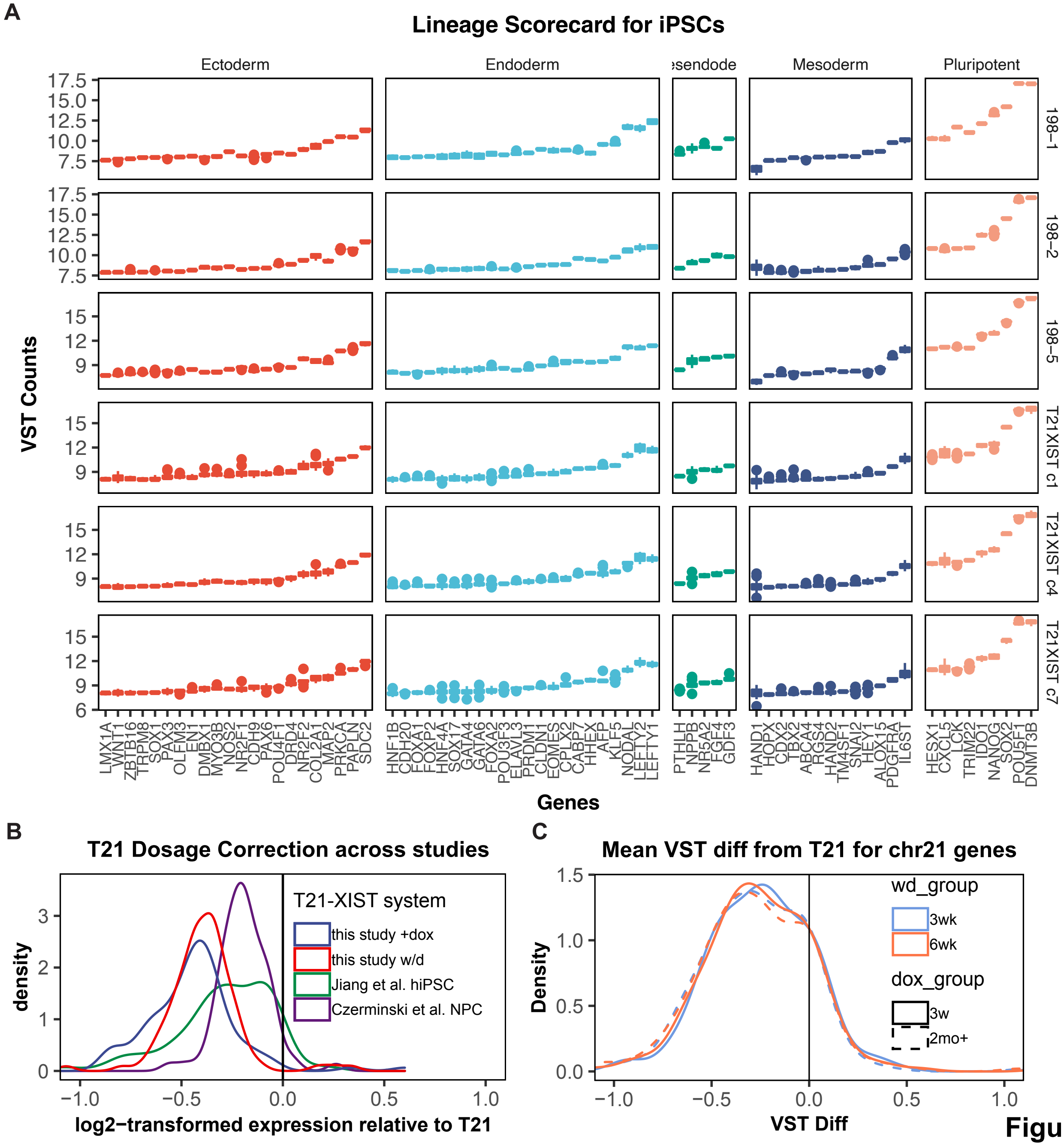
A
Lineage Scorecard for iPSCs
Ectoderm
Endoderm
Mesoderm
Pluripotent
198–1
198–2
198–5
T21XIST c1
T21XIST c4
T21XIST c7
VST Counts
Genes
LMX1A
WNT1
ZBTB16
TRPM8
SOX1
PAX3
OLFM3
EN1
DMBX1
MYO3B
NOS2
NR2F1
CDH9
PAX6
POU4F1
DRD4
NR2F2
COL2A1
MAP2
PRKCA
PAPLN
SDC2
HNF1B
CDH20
FOXA1
FOXP2
HNF4A
SOX17
GATA4
GATA6
FOXA2
POU3F3
ELAVL3
PRDM1
CLDN1
EOMES
CPLX2
CABP7
HHEX
AFP
KLF5
NODAL
LEFTY2
LEFTY1
PTHLH
NPPB
NR5A2
FGF4
GDF3
HAND1
HOPX
CDX2
TBX3
ABCA4
RGS4
HAND2
TM4SF1
SNAI2
HEY1
ALOX15
PDGFRA
IL6ST
HESX1
CXCL5
LCK
TRIM22
IDO1
NANOG
SOX2
POU5F1
DNMT3B
B
T21 Dosage Correction across studies
T21-XIST system
this study +dox
this study w/d
Jiang et al. hiPSC
Czerminski et al. NPC
density
log2−transformed expression relative to T21
C
Mean VST diff from T21 for chr21 genes
wd_group
3wk
6wk
dox_group
3w
2mo+
Density
VST Diff

Figure S2

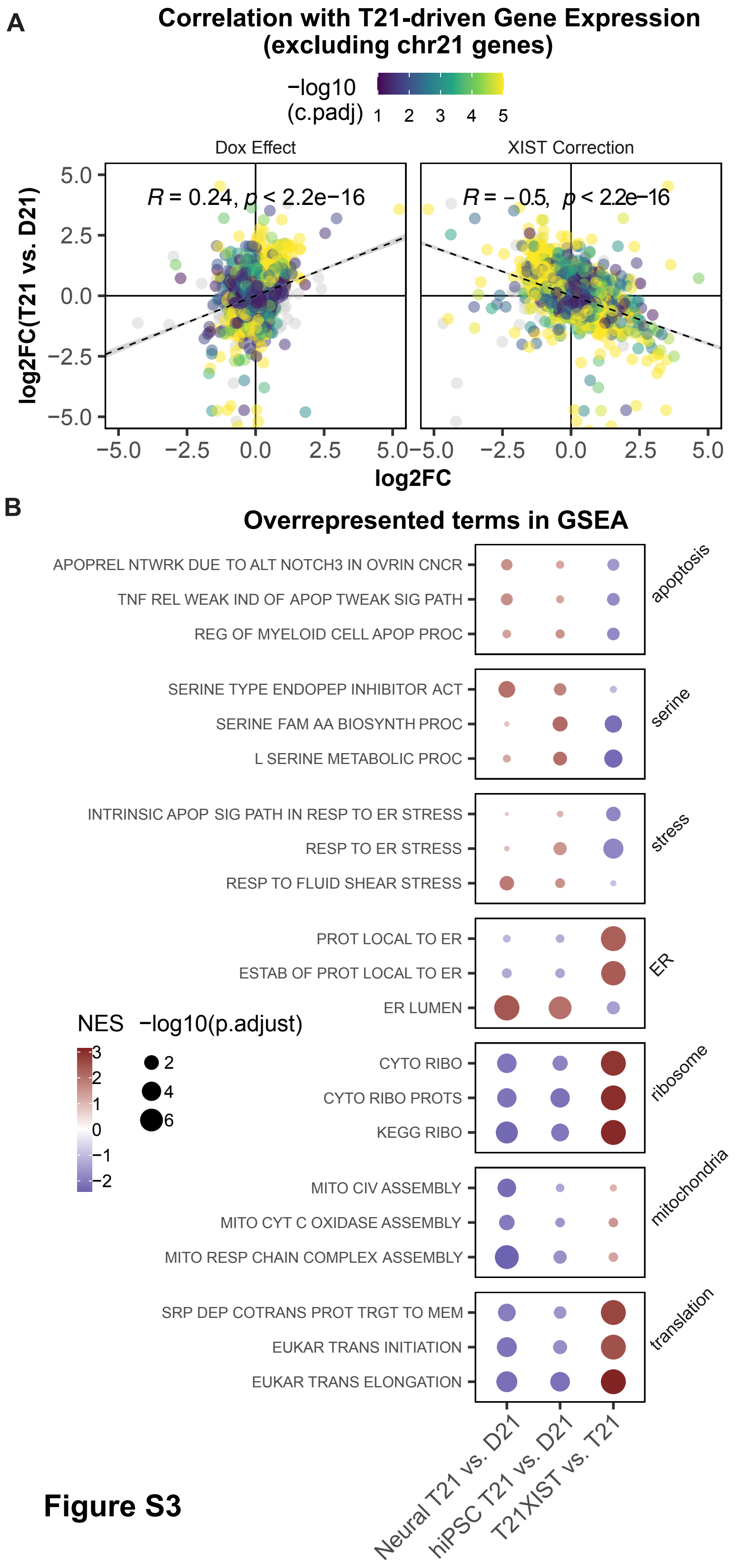
A
Correlation with T21-driven Gene Expression
(excluding chr21 genes)
−log10
(c.padj)
1 2 3 4 5
Dox Effect
XIST Correction
R = 0.24, p < 2.2e−16
R = −0.5, p < 2.2e−16
log2FC(T21 vs. D21)
5.0
2.5
0.0
−2.5
−5.0
−5.0 −2.5 0.0 2.5 5.0
log2FC
B
Overrepresented terms in GSEA
APOPREL NTWRK DUE TO ALT NOTCH3 IN OVRIN CNCR
TNF REL WEAK IND OF APOP TWEAK SIG PATH
REG OF MYELOID CELL APOP PROC
apoptosis
SERINE TYPE ENDOPEP INHIBITOR ACT
SERINE FAM AA BIOSYNTH PROC
L SERINE METABOLIC PROC
serine
INTRINSIC APOP SIG PATH IN RESP TO ER STRESS
RESP TO ER STRESS
RESP TO FLUID SHEAR STRESS
stress
PROT LOCAL TO ER
ESTAB OF PROT LOCAL TO ER
ER LUMEN
ER
CYTO RIBO
CYTO RIBO PROTS
KEGG RIBO
ribosome
MITO CIV ASSEMBLY
MITO CYT C OXIDASE ASSEMBLY
MITO RESP CHAIN COMPLEX ASSEMBLY
mitochondria
SRP DEP COTRANS PROT TRGT TO MEM
EUKAR TRANS INITIATION
EUKAR TRANS ELONGATION
translation
NES
3
2
1
0
−1
−2
−log10(p.adjust)
2
4
6
Neural T21 vs. D21
hiPSC T21 vs. D21
T21XIST vs. T21

**Figure S3**

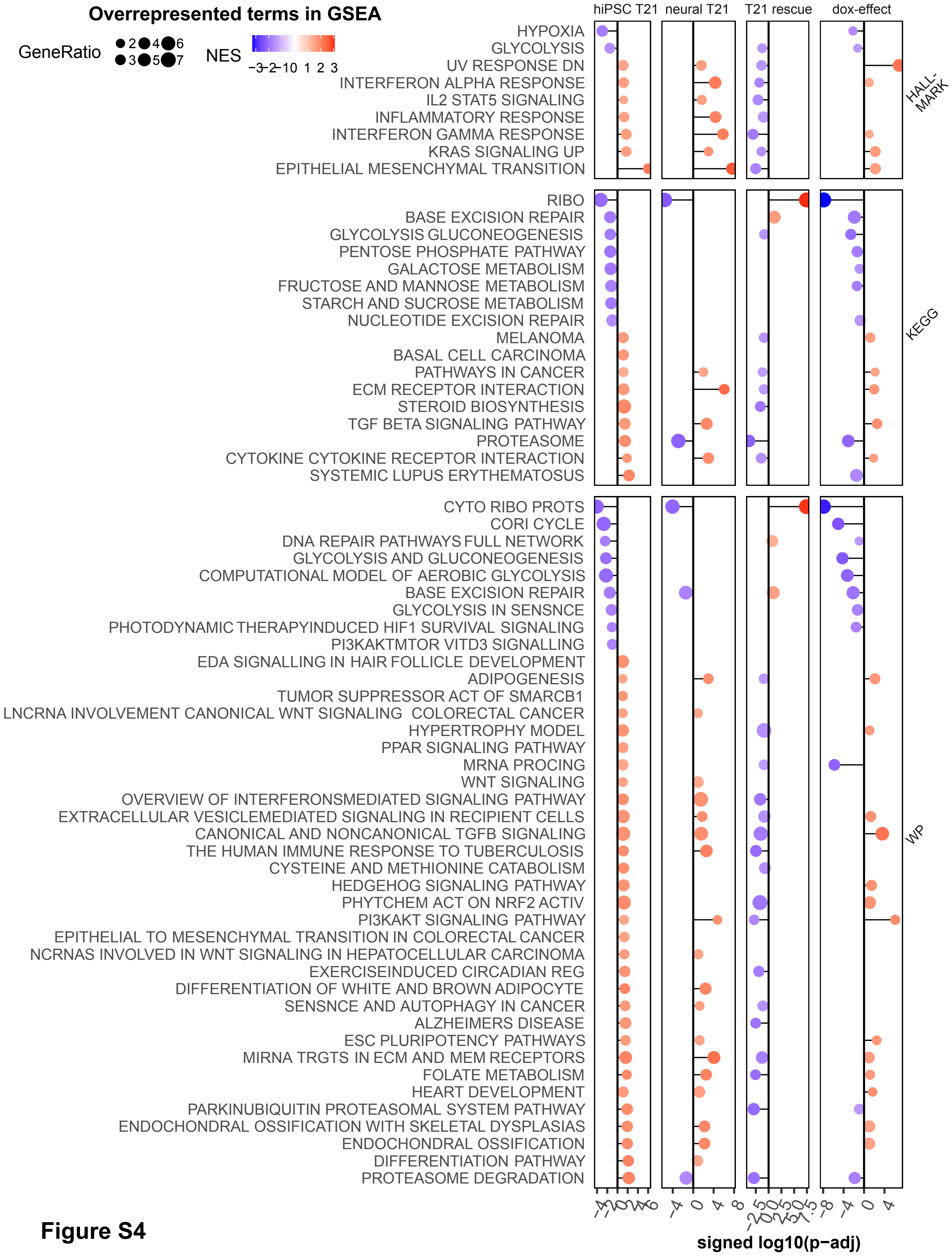
Overrepresented terms in GSEA
GeneRatio
2
3
4
5
6
7
NES
−3
−2
−1
0
1
2
3
hiPSC T21
neural T21
T21 rescue
dox-effect
HYPOXIA
GLYCOLYSIS
UV RESPONSE DN
INTERFERON ALPHA RESPONSE
IL2 STAT5 SIGNALING
INFLAMMATORY RESPONSE
INTERFERON GAMMA RESPONSE
KRAS SIGNALING UP
EPITHELIAL MESENCHYMAL TRANSITION
HALL-MARK
RIBO
BASE EXCISION REPAIR
GLYCOLYSIS GLUCONEOGENESIS
PENTOSE PHOSPHATE PATHWAY
GALACTOSE METABOLISM
FRUCTOSE AND MANNOSE METABOLISM
STARCH AND SUCROSE METABOLISM
NUCLEOTIDE EXCISION REPAIR
MELANOMA
BASAL CELL CARCINOMA
PATHWAYS IN CANCER
ECM RECEPTOR INTERACTION
STEROID BIOSYNTHESIS
TGF BETA SIGNALING PATHWAY
PROTEASOME
CYTOKINE CYTOKINE RECEPTOR INTERACTION
SYSTEMIC LUPUS ERYTHEMATOSUS
KEGG
CYTO RIBO PROTS
CORI CYCLE
DNA REPAIR PATHWAYS FULL NETWORK
GLYCOLYSIS AND GLUCONEOGENESIS
COMPUTATIONAL MODEL OF AEROBIC GLYCOLYSIS
BASE EXCISION REPAIR
GLYCOLYSIS IN SENSNCE
PHOTODYNAMIC THERAPYINDUCED HIF1 SURVIVAL SIGNALING
PI3KAKTMTOR VITD3 SIGNALLING
EDA SIGNALLING IN HAIR FOLLICLE DEVELOPMENT
ADIPOGENESIS
TUMOR SUPPRESSOR ACT OF SMARCB1
LNCRNA INVOLVEMENT CANONICAL WNT SIGNALING COLORECTAL CANCER
HYPERTROPHY MODEL
PPAR SIGNALING PATHWAY
MRNA PROCING
WNT SIGNALING
OVERVIEW OF INTERFERONSMEDIATED SIGNALING PATHWAY
EXTRACELLULAR VESICLEMEDIATED SIGNALING IN RECIPIENT CELLS
CANONICAL AND NONCANONICAL TGFB SIGNALING
THE HUMAN IMMUNE RESPONSE TO TUBERCULOSIS
CYSTEINE AND METHIONINE CATABOLISM
HEDGEHOG SIGNALING PATHWAY
PHYTCHEM ACT ON NRF2 ACTIV
PI3KAKT SIGNALING PATHWAY
EPITHELIAL TO MESENCHYMAL TRANSITION IN COLORECTAL CANCER
NCRNAS INVOLVED IN WNT SIGNALING IN HEPATOCELLULAR CARCINOMA
EXERCISEINDUCED CIRCADIAN REG
DIFFERENTIATION OF WHITE AND BROWN ADIPOCYTE
SENSNCE AND AUTOPHAGY IN CANCER
ALZHEIMERS DISEASE
ESC PLURIPOTENCY PATHWAYS
MIRNA TRGTS IN ECM AND MEM RECEPTORS
FOLATE METABOLISM
HEART DEVELOPMENT
PARKINUBIQUITIN PROTEASOMAL SYSTEM PATHWAY
ENDOCHONDRAL OSSIFICATION WITH SKELETAL DYSPLASIAS
ENDOCHONDRAL OSSIFICATION
DIFFERENTIATION PATHWAY
PROTEASOME DEGRADATION
WP
−4
0
4
8
−8
−4
0
4
signed log10(p−adj)

**Figure S4**

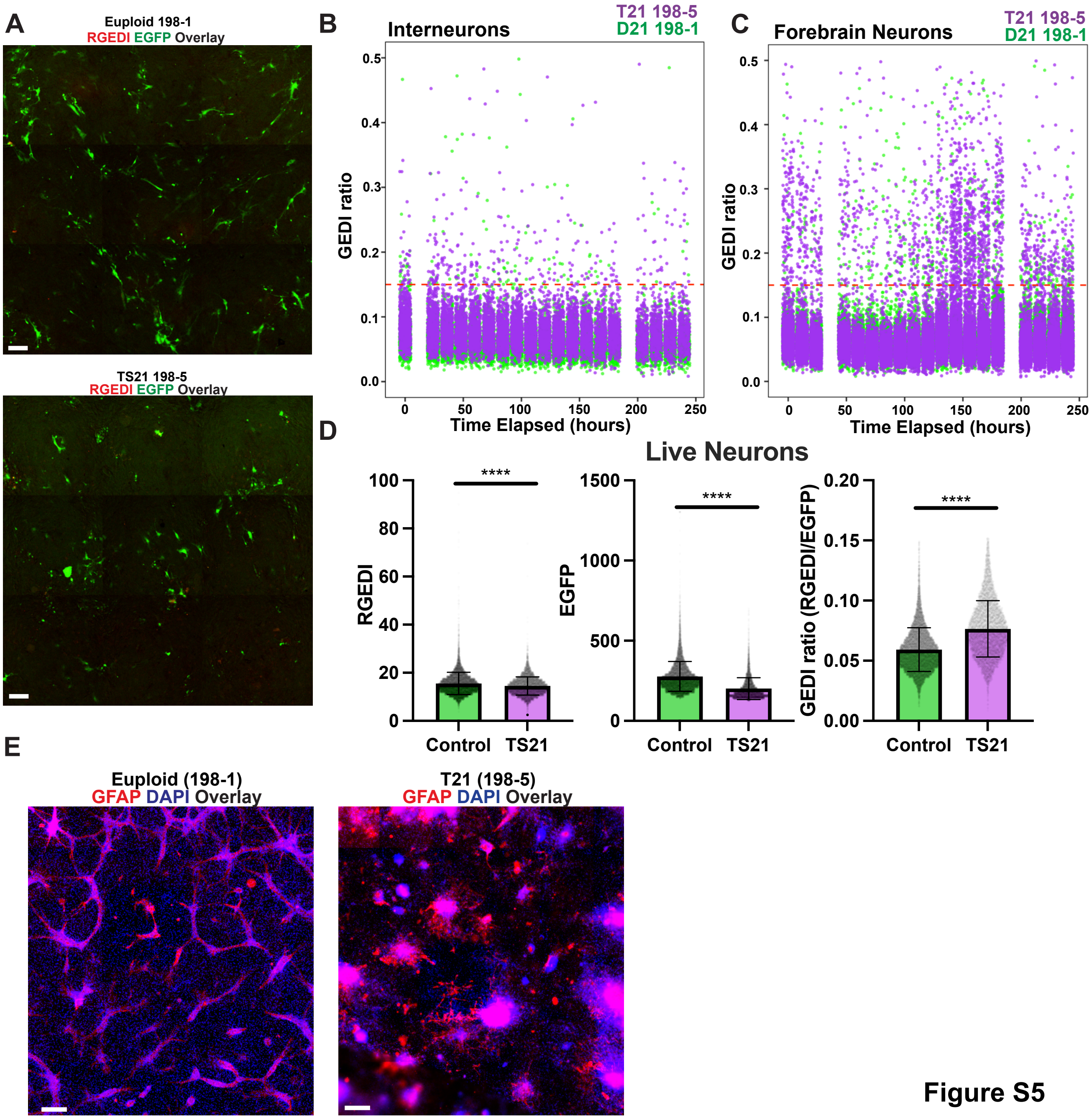
A
Euploid 198-1
RGEDI EGFP Overlay
TS21 198-5
RGEDI EGFP Overlay
B
Interneurons
T21 198-5
D21 198-1
GEDI ratio
Time Elapsed (hours)
C
Forebrain Neurons
T21 198-5
D21 198-1
GEDI ratio
Time Elapsed (hours)
D
Live Neurons
RGEDI
EGFP
GEDI ratio (RGEDI/EGFP)
****
Control
TS21
E
Euploid (198-1)
GFAP DAPI Overlay
T21 (198-5)
GFAP DAPI Overlay

Figure S5

A

B

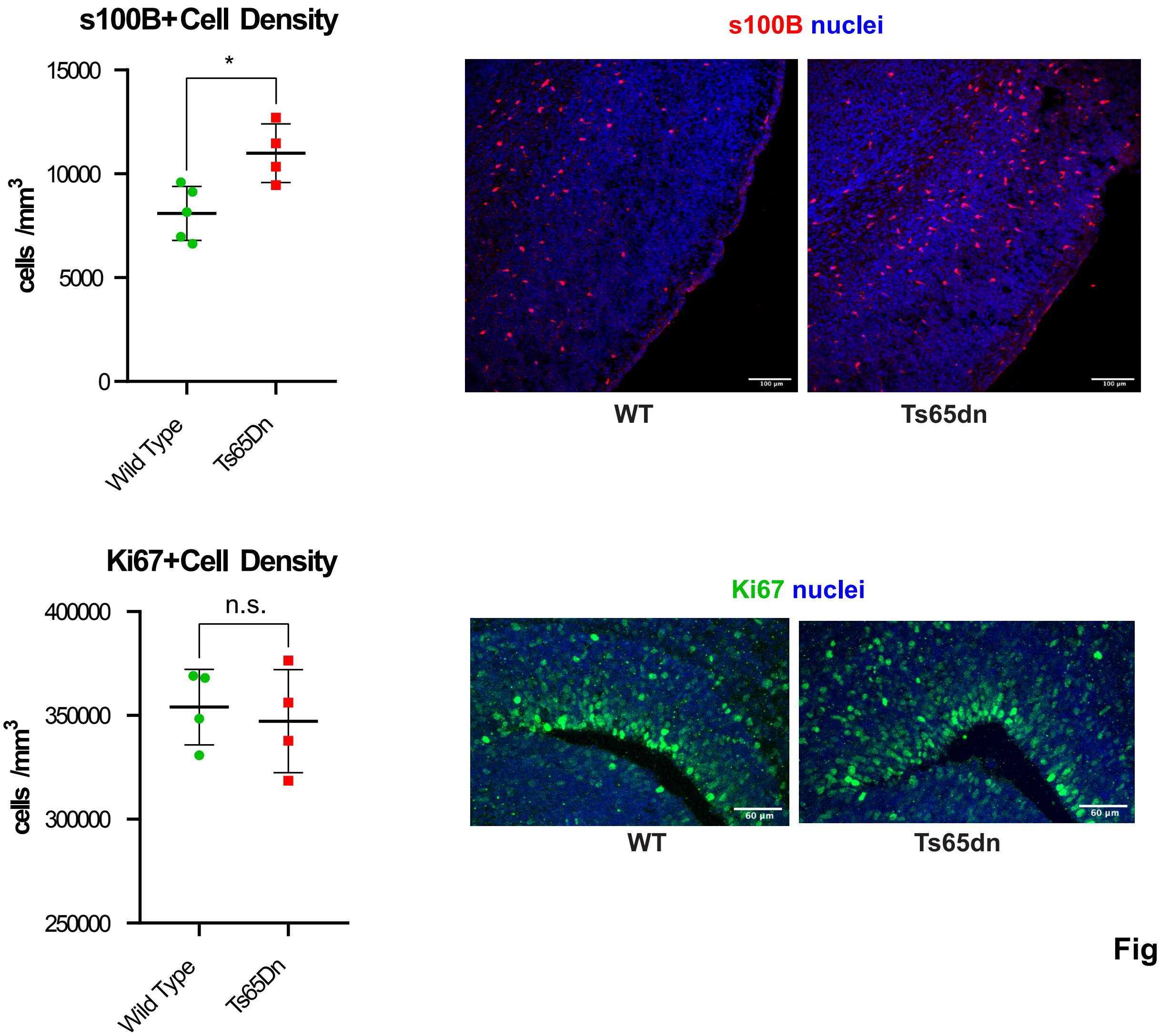
s100B+Cell Density
*
15000
10000
5000
0
cells /mm3
Wild Type
Ts65Dn
s100B nuclei
WT
Ts65dn
Ki67+Cell Density
n.s.
400000
350000
300000
250000
cells /mm3
Wild Type
Ts65Dn
Ki67 nuclei
WT
Ts65dn

Figure S6

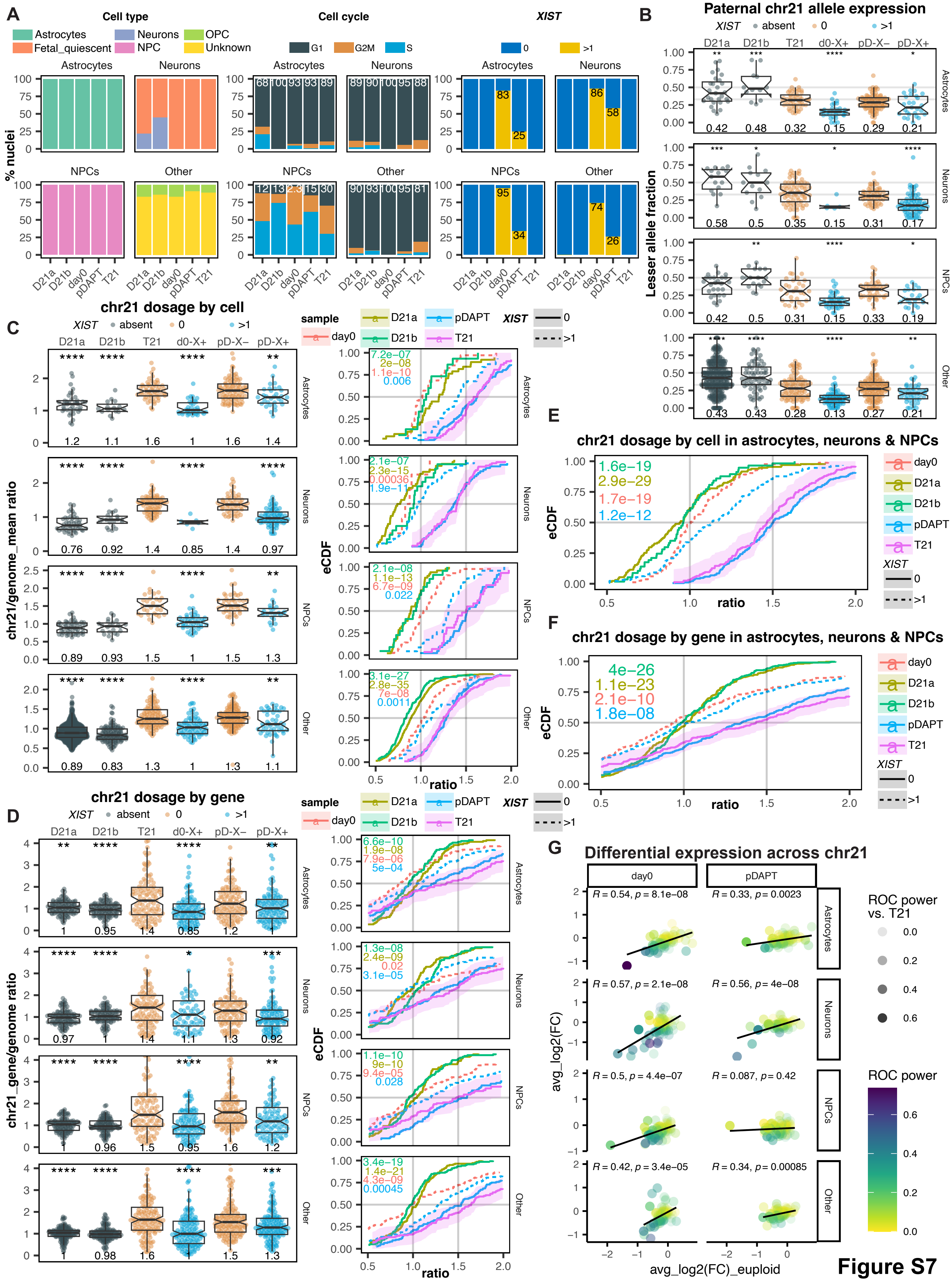

A
Cell type
Astrocytes
Neurons
OPC
Fetal_quiescent
NPC
Unknown
Cell cycle
G1
G2M
S
XIST
0
>1
% nuclei
NPCs
Other
D21a
D21b
day0
pDAPT
T21
B
Paternal chr21 allele expression
absent
d0-X+
pD-X−
pD-X+
Lesser allele fraction
C
chr21 dosage by cell
sample
chr21/genome_mean ratio
eCDF
ratio
D
chr21 dosage by gene
chr21_gene/genome ratio
E
chr21 dosage by cell in astrocytes, neurons & NPCs
F
chr21 dosage by gene in astrocytes, neurons & NPCs
G
Differential expression across chr21
ROC power vs. T21
ROC power
avg_log2(FC)
avg_log2(FC)_euploid


Figure S7

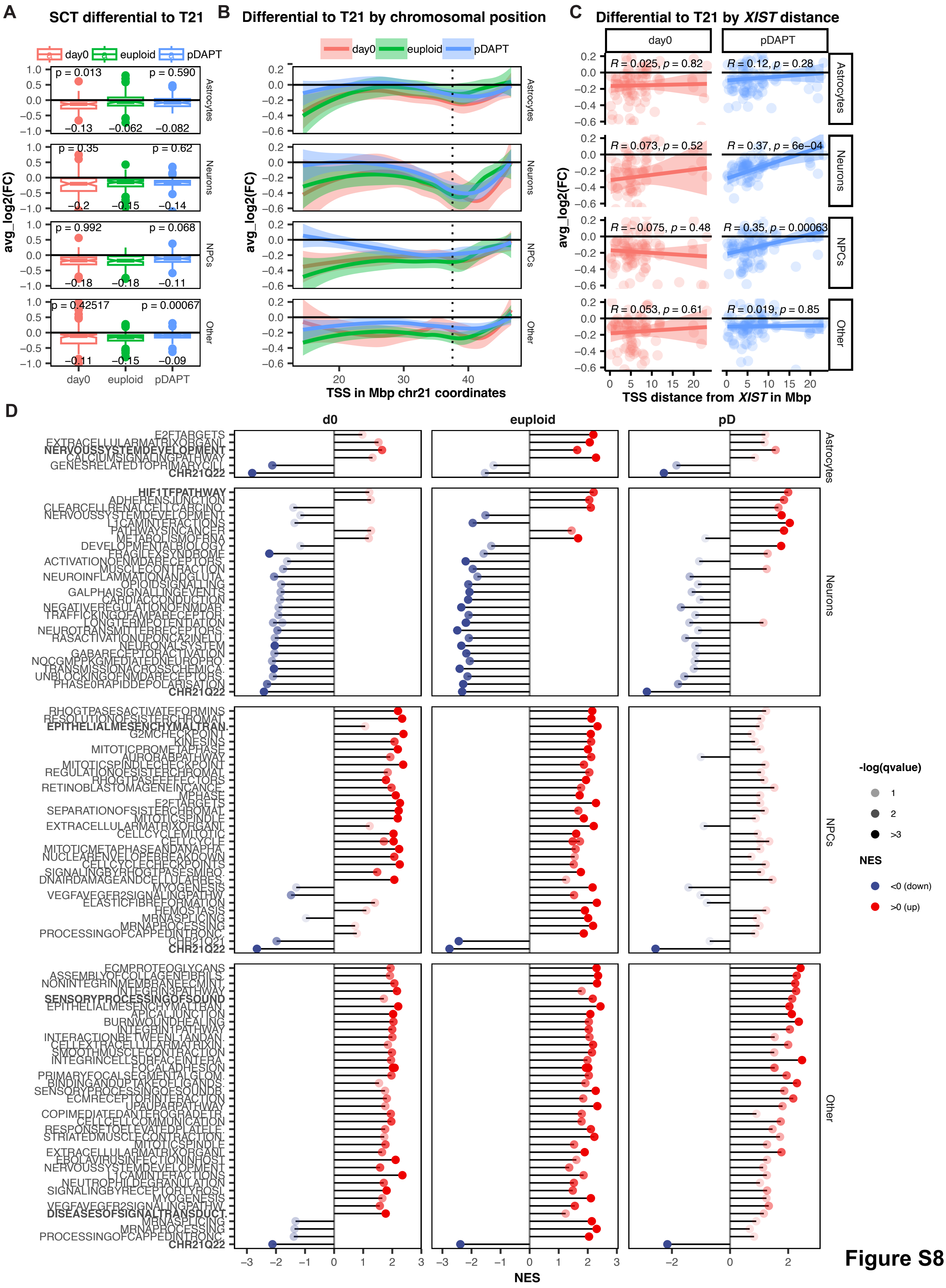

A
SCT differential to T21
day0
euploid
pDAPT
p = 0.013
p = 0.590
−0.13
−0.062
−0.082
p = 0.35
p = 0.62
−0.2
−0.15
−0.14
p = 0.992
p = 0.068
−0.18
−0.18
−0.11
p = 0.42517
p = 0.00067
−0.11
−0.15
−0.09
avg_log2(FC)
Astrocytes
Neurons
NPCs
Other
B
Differential to T21 by chromosomal position
TSS in Mbp chr21 coordinates
C
Differential to T21 by XIST distance
R = 0.025, p = 0.82
R = 0.12, p = 0.28
R = 0.073, p = 0.52
R = 0.37, p = 6e−04
R = −0.075, p = 0.48
R = 0.35, p = 0.00063
R = 0.053, p = 0.61
R = 0.019, p = 0.85
TSS distance from XIST in Mbp
D
d0
euploid
pD
E2FTARGETS
EXTRACELLULARMATRIXORGANI.
NERVOUSSYSTEMDEVELOPMENT
CALCIUMSIGNALINGPATHWAY
GENESRELATEDTOPRIMARYCILI.
CHR21Q22
HIF1TFPATHWAY
ADHERENSJUNCTION
CLEARCELLRENALCELLCARCINO.
NERVOUSSYSTEMDEVELOPMENT
L1CAMINTERACTIONS
PATHWAYSINCANCER
METABOLISMOFRNA
DEVELOPMENTALBIOLOGY
FRAGILEXSYNDROME
ACTIVATIONOFNMDARECEPTORS.
MUSCLECONTRACTION
NEUROINFLAMMATIONANDGLUTA.
OPIOIDSIGNALLING
GALPHAISIGNALLINGEVENTS
CARDIACCONDUCTION
NEGATIVEREGULATIONOFNMDAR.
TRAFFICKINGOFAMPARECEPTOR.
LONGTERMPOTENTIATION
NEUROTRANSMITTERRECEPTORS.
RASACTIVATIONUPONCA2INFLU.
NEURONALSYSTEM
GABARECEPTORACTIVATION
NOCGMPPKGMEDIATEDNEUROPRO.
TRANSMISSIONACROSSCHEMICA.
UNBLOCKINGOFNMDARECEPTORS.
PHASE0RAPIDDEPOLARISATION
CHR21Q22
RHOGTPASESACTIVATEFORMINS
RESOLUTIONOFSISTERCHROMAT.
EPITHELIALMESENCHYMALTRAN.
G2MCHECKPOINT
KINESINS
MITOTICPROMETAPHASE
AURORABPATHWAY
MITOTICSPINDLECHECKPOINT
REGULATIONOFSISTERCHROMAT.
RHOGTPASEEFFECTORS
RETINOBLASTOMAGENEINCANCE.
MPHASE
E2FTARGETS
SEPARATIONOFSISTERCHROMAT.
MITOTICSPINDLE
EXTRACELLULARMATRIXORGANI.
CELLCYCLEMITOTIC
CELLCYCLE
MITOTICMETAPHASEANDANAPHA.
NUCLEARENVELOPEBREAKDOWN
CELLCYCLECHECKPOINTS
SIGNALINGBYRHOGTPASESMIRO.
DNAIRDAMAGEANDCELLULARRES.
MYOGENESIS
VEGFAVEGFR2SIGNALINGPATHW.
ELASTICFIBREFORMATION
HEMOSTASIS
MRNASPLICING
MRNAPROCESSING
PROCESSINGOFCAPPEDINTRONC.
CHR21Q21
CHR21Q22
ECMPROTEOGLYCANS
ASSEMBLYOFCOLLAGENFIBRILS.
NONINTEGRINMEMBRANEECMINT.
INTEGRIN3PATHWAY
SENSORYPROCESSINGOFSOUND
EPITHELIALMESENCHYMALTRAN.
APICALJUNCTION
BURNWOUNDHEALING
INTEGRINTPATHWAY
INTERACTIONBETWEENL1ANDAN.
CELLEXTRACELLULARMATRIXIN.
SMOOTHMUSCLECONTRACTION
INTEGRINCELLSURFACEINTERA.
FOCALADHESION
PRIMARYFOCALSEGMENTALGLOM.
BINDINGANDUPTAKEOFLIGANDS.
SENSORYPROCESSINGOFSOUNDB.
ECMRECEPTORINTERACTION
UPAURPATHWAY
COPIMEDIATEDANTEROGRADETR.
CELLCELLCOMMUNICATION
RESPONSETOELEVATEDPLATELE.
STRIATEDMUSCLECONTRACTION
MITOTICSPINDLE
EXTRACELLULARMATRIXORGANI.
EBOLAVIRUSINFECTIONINHOST.
NERVOUSSYSTEMDEVELOPMENT
L1CAMINTERACTIONS
NEUTROPHILDEGRANULATION
SIGNALINGBYRECEPTORTYROSI.
MYOGENESIS
VEGFAVEGFR2SIGNALINGPATHW.
DISEASESOFSIGNALTRANSDUCT.
MRNASPLICING
MRNAPROCESSING
PROCESSINGOFCAPPEDINTRONC.
CHR21Q22
NES
-log(qvalue)
1
2
>3
NES
<0 (down)
>0 (up)


Figure S8
